# Mental fatigue impairs cycling endurance performance and perception of effort, but not muscle activation

**DOI:** 10.64898/2026.03.19.712281

**Authors:** Robin Souron, Aurélie Sarcher, Lilian Lacourpaille, Inès Boulaouche, Calvin Richier, Thomas Mangin, Mathieu Gruet, Julie Doron, Marc Jubeau, Benjamin Pageaux

**Affiliations:** Nantes Université, Movement - Interactions - Performance, MIP, UR 4334, F-44000 Nantes, France; Université de Toulon, Laboratoire IAPS (n°201723207F), Toulon, France; École de kinésiologie et des sciences de l’activité physique (EKSAP), Faculté de Médecine, Université de Montréal, Montréal, Canada; Centre de recherche de l’Institut universitaire de gériatrie de Montréal (CRIUGM), Montréal, Canada; Centre Interdisciplinaire de Recherche sur le Cerveau et l’Apprentissage (CIRCA), Montréal, Canada

**Author notes:** Corresponding author: Robin Souron, Laboratoire « Motricité, Interactions, Performance » (EA 4334) 25 bis, Boulevard Guy Mollet, BP 72206 - 44322 NANTES cedex 3.

**Keywords:** Cognitive exertion, physical performance, motor command, electromyography, perceived exertion

## Abstract

Mental fatigue is induced by prolonged engagement in cognitively demanding tasks and impairs endurance performance. The neuropsychophysiological mechanisms underlying this decreased performance remain unclear, with suggestion that mental fatigue may disrupt motor command and consequently muscle activation. We aimed to test this hypothesis in a repeated cross-over design study in which 18 participants completed two experimental sessions involving a time-to-task failure cycling test at 80% of peak power output. Each cycling task was preceded by 1h of a prolonged Stroop task (Stroop condition) or a neutral control task (Control condition). Mental fatigue was assessed using a visual analog scale anchored with "not fatigued at all" and "extremely fatigued. Perception of effort and surface electromyography from ten lower-limb muscles of the right leg were recorded at regular intervals during cycling. Mental fatigue was higher in the Stroop compared to the Control condition (*p* = .002). Endurance cycling time was shorter in the Stroop than in the Control condition (887 ± 284 s vs. 999 ± 379 s, respectively; −111 ± 160 s, p = .009). No significant differences in electromyography parameters were observed between Stroop and Control conditions, for any muscle (*p* > .05). Perception of effort was higher in the Stroop condition from the onset of the cycling task (*p* = .006), and the rate of increase in perception of effort was significantly higher in the Stroop than Control condition (*p* = .031). Our findings do not support the hypothesis that mental fatigue alters motor control or increases central motor command, as no changes in muscle activation were detected. Conversely, our results reinforce the notion that prolonged cognitive engagement impairs endurance performance primarily through an increased perception of effort. Future research should consider combining surface electromyography with more sensitive neurophysiological techniques to investigate potential subtle changes in motor drive during dynamic, whole-body tasks under mental fatigue.

## Introduction

Mental fatigue is commonly defined as a psychobiological state induced by prolonged engagement in demanding cognitive tasks (Boksem and Tops 2008; Marcora et al. 2009). It should be distinguished from cognitive performance decrement, which may occur as one possible consequence of mental fatigue but is not required for mental fatigue to be present (Pageaux et al. 2013; Pageaux et al. 2015; Mangin and Pageaux 2025). Subjectively, mental fatigue is characterized by an increase in tiredness and lack of energy (Boksem and Tops 2008), and an aversion to continuing or initiating tasks (Müller and Apps 2019). Objectively, mental fatigue alters cognitive functioning, leading to decreased cognitive performance or the maintenance of performance at the cost of increased effort (Mangin and Pageaux 2025). Beyond the negative effect of mental fatigue on cognitive performance, numerous studies have also reported its detrimental effects on physical performance (Van Cutsem et al. 2017; Brown et al. 2020; Pageaux and Lepers 2018; Giboin and Wolff 2019). Mental fatigue has been shown to reduce endurance performance in isometric tasks performed to task failure (Bray et al. 2008; Pageaux et al. 2013; Alix-Fages et al. 2023). The impairment in endurance performance has also been observed during cycling and running endurance tests (Marcora et al. 2009; MacMahon et al. 2014; Smith et al. 2016; Martin et al. 2016). This reduction in endurance performance has been proposed to be associated with a higher-than-normal perception of effort (Marcora et al. 2009; Pageaux and Lepers 2018).

The perception of effort during a physical task, defined as the conscious awareness of how hard, heavy, and strenuous the task feels (Marcora 2010), is a key regulator of endurance performance (Pageaux 2014; Marcora et al. 2008; Marcora and Staiano 2010). At the behavioral level, building on the intensity of motivation theory (Brehm and Self 1989; Richter et al. 2016), the perception of effort can be linked to the regulation of endurance performance through the psychobiological model of endurance performance (Marcora 2019; Marcora et al. 2008; Pageaux 2014). According to this framework, task disengagement, also referred to as exhaustion or task failure, occurs when an individual exerts the maximal effort they are willing to invest in the task. This upper limit is determined by the individual’s motivation to succeed (Marcora 2019; Marcora et al. 2008; Pageaux 2014). Previous studies have demonstrated that, for the same mechanical output, the perception of effort increases following prolonged engagement (≥ 30 min) in cognitive tasks (Marcora et al. 2009; Pageaux et al. 2013). This heightened perception of effort leads individuals to reach the maximum effort they are willing to exert earlier, resulting in task disengagement and impaired endurance performance.

Understanding the neurophysiological alterations that lead to an increased perception of effort in the presence of fatigue could help unravel the origins of the detrimental effects of mental fatigue on endurance performance. The perception of effort is generated by the brain’s processing of sensory signals. Although the nature of these signals remains debated, converging evidence suggests that effort perception originates from signals related to the motor command (for reviews, see Marcora (2009); Bergevin et al. (2023); Mangin and Pageaux (2026)). During voluntary muscle contractions, motor commands are sent to the working muscles, and an efference copy of these motor commands, referred to as the corollary discharge, is transmitted to cortical areas involved in sensory processing (Christensen et al. 2007; McCloskey 1981). The corollary discharge model proposes that perceived effort is generated through the brain’s processing of this signal (De Morree et al. 2012; Bergevin et al. 2023). Within this framework, an alteration in perceived effort during physical tasks may result from changes in motor commands and their corollary discharges, from altered brain processing of these signals, or from both.

In line with this model, the increase in perceived effort under mental fatigue may be linked to specific modulations of (pre)frontal cortex activity, potentially driven by metabolic alterations (Barakat et al. 2025; Pessiglione et al. 2025). Among frontal brain areas, the anterior cingulate cortex is likely a key structure in the negative effects of mental fatigue on the perception of effort, and consequently, on endurance performance. The anterior cingulate cortex plays a crucial role in cognitive control and effort-based decision-making (Shenhav et al. 2016), and its activity increases during demanding cognitive tasks (Carter et al. 1998; Pardo et al. 1990). The anterior cingulate cortex is proposed to interact with the prefrontal cortex (Hogan et al. 2019; Paus 2001), another key area involved in the regulation of cognitive performance and decision-making (Domenech and Koechlin 2015), proposed to be involved in task disengagement (Meeusen et al. 2016; Robertson and Marino 2016) and altered by the presence of mental fatigue (Yan et al. 2025). Specific cingulate regions project extensively to premotor and motor cortices, as well as to the spinal cord, all key structures involved in motor command and control (Medford and Critchley 2010; Paus 2001; Gillies et al. 2019). These connections suggest that cognitively demanding tasks inducing mental fatigue could disrupt motor command by altering prefrontal circuitry involved in the regulation of the motor command during task execution. Such interactions may influence muscle activation and could explain reports of increased muscle activity under mental fatigue during isometric exercise (Bray et al. 2008; Jacquet et al. 2021). However, a recent study observed that mental fatigue induced by a prolonged Stroop task, while decreasing dorsiflexion isometric endurance performance, did not affect motor command and muscle fibers recruitment assessed with high density-electromyography (EMG) (Alix-Fages et al. 2023). The discrepancies between these studies highlight the need to further investigate how mental fatigue influences muscle activation, particularly during dynamic exercise modalities, such as cycling, which better reflect real-world exercise condition. Indeed, isometric contractions may not fully capture the complexity of motor control during ecological movement. In this context, increased vastus lateralis activity has been observed during a submaximal cycling task under mental fatigue (Pageaux et al. 2015). However, because cycling requires the coordinated activation of multiple muscles (So et al. 2005), the results of Pageaux et al. (2015), which focused on a single muscle, do not provide a comprehensive understanding of the effects of mental fatigue on muscle activation and should therefore be replicated using a more comprehensive assessment of lower-limb muscle activation.

The present study aimed to investigate whether the presence of mental fatigue is associated with alterations in muscle activation during dynamic exercise. In secondary analysis, we aimed to replicate the negative effect of mental fatigue on endurance performance. We selected cycling as the endurance exercise because it allows for a more precise and comprehensive analysis of lower-limb muscle EMG activity compared to running, as previously performed in our laboratory (Hug et al. 2019). We hypothesized that mental fatigue would increase perception of effort and alter lower-limb muscle activation, thereby reducing endurance performance.

## Methods

### Study design and overview of the laboratory visits

This study used a within-subject design, with participants visiting the laboratory on three separate occasions. The first visit was for familiarization purpose, and the second and third visits were the control and experimental conditions (randomized, counterbalanced order). All visits were spaced at least 72 h apart, with a maximum interval of 7 days between the two experimental conditions. The control and experimental conditions were identical, except for the cognitive task performed: watching a documentary (Control condition) or performing a Stroop task (Stroop condition). Each session began with questionnaires about mental fatigue and motivation. EMG electrodes were then placed on ten muscles of the right leg and signal quality was verified (see the *“Electromyography”* section). Participants then engaged either in 1 h of the Stroop task or 1 h of viewing an emotionally neutral documentary. Mental fatigue, motivation and boredom were measured during and post Stroop and documentary. Then, participants promptly transferred to the cycle ergometer and performed a cycling time to task failure test at 80% of their peak power output (PPO) until task failure.

### Participants

Twenty participants (8 female; age: 22 ± 5 years; height: 175 ± 12 cm; weight: 68 ± 14 kg; <u>peak power</u> output: 250 ± 65 W; 3.7 ± 0.6 W.kg^-1^) participated in this repeated cross-over design study. Participants reported engaging in 2-6 h of moderate physical activity per week, but did not report extensive cycling training history and were not currently active cyclists. Participants were instructed to refrain from engaging in any strenuous lower-limb physical activity during the 24 h preceding each laboratory visit. Participants were provided with written instructions detailing all study procedures but were unaware of its objectives and hypotheses. They were not informed that the cognitive task was used to induce mental fatigue and modify the subsequent endurance performance. Written informed consent was obtained from each participant. This study adhered to the standards outlined in the latest revision of the Declaration of Helsinki (with the exception of registration in a database) and was approved by the Nantes University ethics committee (n° IRB: IORG0011023).

The sample size rationale was first based on the aim of matching the sample size of the first influential study on the effects of mental fatigue on endurance performance, which included 16 participants (Marcora et al., 2009), while accounting for potential attrition. Second, although we acknowledge the need to increase sample sizes in mental fatigue research to better estimate the true effect of mental fatigue on endurance performance, our sample size was constrained by the time resources available for the MSc students recruited as experimenters in this study. We therefore recruited as much participants as possible in this given time (Lakens 2022). We recruited 20 participants to secure a final sample size at least comparable to this previous work. Two participants were excluded after reporting engagement in strenuous physical activity a few hours prior to the experimental testing. As a result, 18 participants were included in the statistical analysis. A sensitivity analysis revealed that with our sample size (n = 18), the minimal detectable effect size (Cohen’s d) for a paired t-test is 0.70 with 80% power and α = 0.05. Regarding our Statistical Parametric Mapping (SPM) analysis, as it includes a correction for temporal smoothness and multiple comparisons across the time series, the actual minimal detectable effect size is likely higher. Thus, only medium-to-large effects could be reliably detected in our study.

### Visit 1: Familiarization and incremental test

Participants began by determining their preferred pedaling cadence on a cycle ergometer set up in hyperbolic mode (Excalibur Sport, Lode, Groningen, Netherlands). Participants cycled for 1 min at 100 W, followed by 1 min at 175 W, and 1 min at 250 W. Participants were instructed to cycle at a freely chosen cadence between 60–90 rpm, and this preferred cadence was recorded for each stage. If a participant did not report at least a moderate effort when cycling at 250W, additional 1-min bouts with increments of 50 W were performed until the participant had to produce a moderate effort (23 out of 100 on the CR100 Borg scale). The preferred cadence recorded at the power output closest to 80% of the participant’s PPO was used during the time to task failure test performed in the Control and Stroop conditions (72 ± 6 rpm).

After a 5-min rest, participants performed an incremental test at their predetermined preferred cadence, starting at 100 W with increments of 25 W every minute until task failure. Task failure was defined as either voluntary disengagement from the test or the inability to maintain the target cadence within ± 3 rpm for more than 3 s. The last fully completed stage was used to determine the participant’s PPO. This PPO, was used to determine the workload for the time to task failure tests performed in the Control and Stroop conditions. Participants were then familiarized with the Stroop task. If participants did not clearly demonstrate an understanding of the instructions, based on experimenter observation and self-reported understanding and comfort with the task, they completed additional familiarization blocks until the task was fully grasped. Then, participants were provided with detailed instructions on how to dissociate effort and muscle pain perceptions, and how to report the intensity of each perception with the CR100 scale (Borg and Kaijser 2006; Pageaux et al. 2020). Finally, participants performed the cycling time to task failure test at 80% PPO for familiarization.

### Visit 2 and 3: Stroop and Control conditions

Participants began each session by engaging, in a randomized and counterbalanced order, for 60- min in an experimental (Stroop task) or control (watched a documentary) condition. In the control and Stroop conditions, mental fatigue and motivation to perform the cycling task were assessed before, at 30- min (mid), and immediately post cognitive tasks. Boredom was also measured mid and post cognitive tasks. Upon completing the questionnaires, participants transferred to the cycle ergometer to begin the cycling time to task failure test. Mental fatigue and motivation to perform the cycling time to task failure test was also measured before and after the test. An overview of the Control and Stroop experimental sessions is presented in Figure 1.

**Figure 1.**
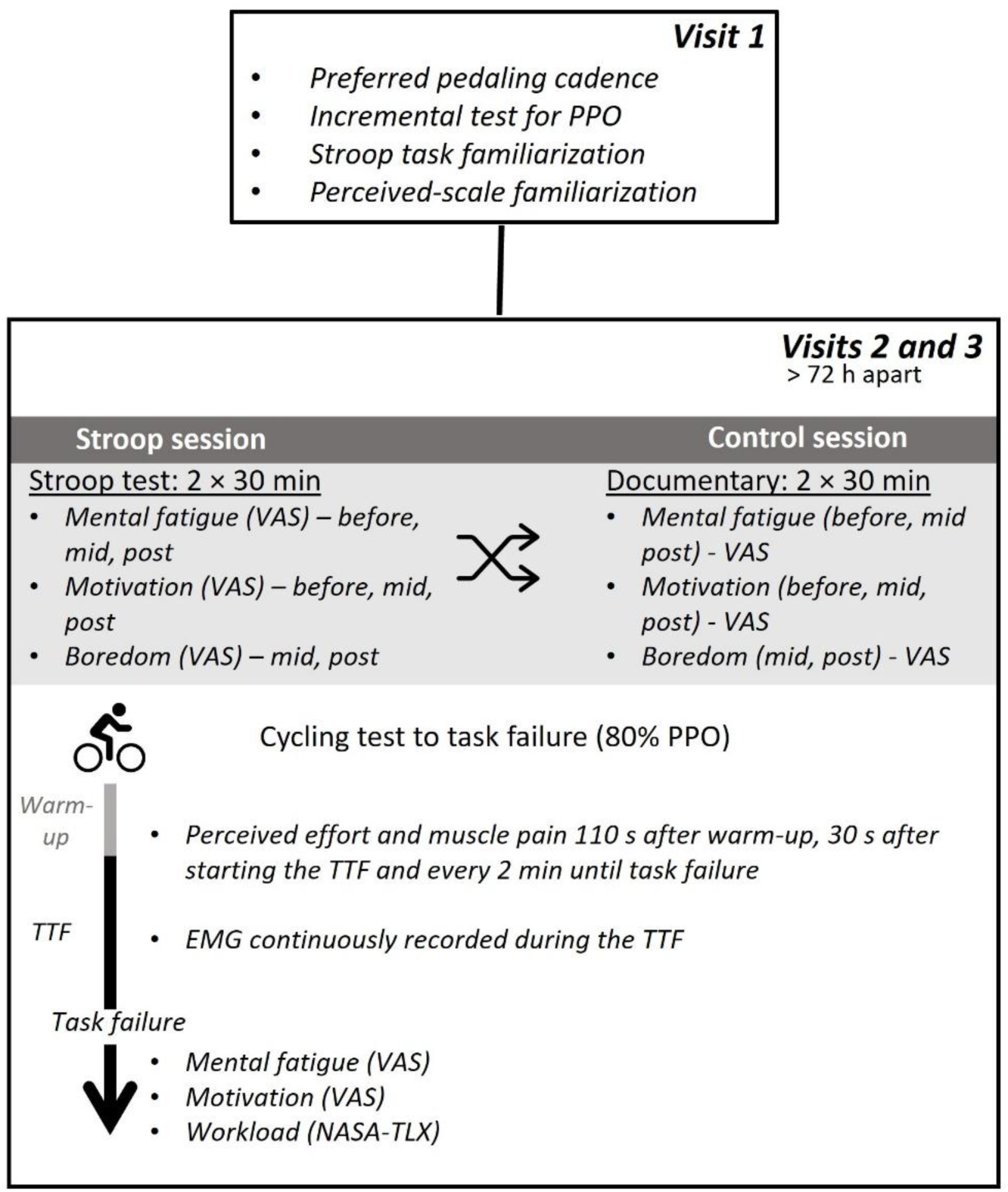
Overview of the experimental and control sessions. Participants visited the laboratory on three distinct occasions. The first visit was dedicated to peak power output (PPO) measurement and familiarization procedures. The second and third visits were the experimental conditions. After completing either a one-hour Stroop task (Stroop condition; two 30-min blocks) or watching a documentary (control condition), participants performed a 3-min warm-up at 40% PPO, followed by a time to task failure test at 80% PPO. Participants reported their perceived effort and muscle pain at regular intervals during the cycling test. Electromyographic (EMG) activity was continuously recorded and analyzed over the 20 pedaling cycles preceding each effort and pain perception measurement. We used visual analogue scales to measure mental changes in mental fatigue, motivation and boredom during the conditions. Participants completed the NASA-TLX at the very end of the procedure.

### Stroop task

Participants performed a modified version of the incongruent Stroop task (Mangin et al. 2021). This task was selected as it represents one of the most widely used cognitive paradigms to induce mental fatigue in the context of physical performance research (Pageaux and Lepers 2018; Schampheleer et al. 2025). The task was performed while seated in an isolated room, using the SuperLab® software (Cedrus, USA), displayed on a 16-inch monitor placed on a desk directly in front of the participant. In visit 1, participants familiarized with the Strop task by completing two blocks of 24 words. In the experimental condition, participants completed the incongruent Stroop task for 1 h, with two bouts of 30-min separated by a 1-min period for mental fatigue, motivation and boredom measurement. Each trial began with the presentation of a fixation cross point enclosed in either a square or a circle for 50 ms at the center of the screen that was positioned in front of the participant. The fixation cross alone remained in the middle of the screen for 400 ms. Immediately after this preparatory signal, a word, the response signal, appeared at the center of the screen and remained visible until the participant responded orally. In cases of omission, the response signal was displayed for 1250 ms before being replaced by a fixation cross for 300 ms. If a participant responded before the end of the 1250 ms, a fixation cross appeared at the center of the screen and remained visible for the remainder of the 1250 ms, plus an additional 300 ms. As a result, each trial lasted exactly 2 seconds. The response signals consisted of color names (i.e., red, blue, yellow, or green) written in an incongruent ink color (e.g., red written in blue). In this task, the meaning of the word was always incongruent with the ink color used to display it. When the fixation cross was enclosed in a square, participants were instructed to read the word, whereas when it was enclosed in a circle, they were required to name the color of the ink. In 50% of the trials, participants were required to read the word, while in the remaining 50%, they had to name the ink color. This 50/50 distribution was chosen to minimize learning effects during the incongruent Stroop task (Dulaney and Rogers 1994), and to increase task difficulty by engaging cognitive flexibility in addition to inhibitory control. The order of trial presentation was randomized.

### Documentary watching

Each participant watched the French version of the documentary *Home* by Y.A. Bertrand. This documentary has been used as an appropriate control condition in the context of mental fatigue studies (Jacquet et al. 2021). The duration of this control task was matched to that of the Stroop task. Participants sat in an isolated room, where the documentary was projected onto a screen positioned 1 m in front of them.

### Cycling time to task failure test

After completion of the cognitive tasks and following a 3-min warm-up at 40% PPO, participants completed a time to task failure test at 80% PPO at their preferred cadence determined in visit 1. This endurance test and intensity were selected to allow direct comparison with previous studies that investigated the effects of mental fatigue on cycling endurance performance using similar protocols (Marcora 2009; Azevedo et al. 2016; Blanchfield et al. 2014). Participants had to remain seated on the saddle. To prevent changes in hip angle caused by lowering the hands on the handlebars, participants were instructed to keep their hands in the same position throughout the test (i.e., on top of the handlebars). Both instructions were visually controlled by the experimenters. Participants reported the intensity of perceived effort and muscle pain every 2 min during the task. No encouragement was provided at any point during the time to task failure test.

Endurance performance was quantified as time to task failure, defined as the time at which participants voluntarily disengaged from the task or were unable to maintain their preferred cadence (± 3 rpm) for more than 3 s despite receiving a standardized, neutral, verbal reminder to return to the target cadence. To compare EMG and perceptual responses during the Stroop and Control conditions, we used individual isotime analysis (Nicolò et al. 2019). For each participant, their shortest time to task failure across the Stroop and Control conditions was defined as 100% isotime. Two other time points, corresponding to 0% and 50% of this 100% isotime, were then calculated for each participant and condition. The 0% isotime time point was always the first measurement available during the time to task failure test.

### EMG

#### EMG recording and electrode placement

Surface EMG was continuously recorded at 2000 Hz during the cycling exercise using a wireless EMG system (Wave Plus wireless system, Cometa, Bareggio MI, Italy) and dedicated software (WavePlus 1.4.0, Cometa, Bareggio MI, Italy). Data were collected for ten muscles of the right lower limb: gluteus maximus (GMax), semitendinous (ST), biceps femoris (BF), vastus medialis (VM), vastus lateralis (VL), rectus femoris (RF), gastrocnemius medialis (GM), gastrocnemius lateralis (GL), soleus (SOL) and tibialis anterior (TA). A pair of self-adhesive Ag/AgCl electrodes (Covidien, Mansfield, MA, USA) was placed on each muscle with a 20-mm center-to-center inter-electrode distance. Electrode placements followed SENIAM recommendations, with the skin shaved and cleaned with alcohol to minimize impedance. To prevent movement-induced artifacts, the wireless EMG units were secured with adhesive tape. The electrode positions established during visit 2 was marked to ensure consistent placement in visit 3.

### Data analysis

Custom-written Matlab (Mathworks, Natick, USA) routines were used to process EMG data, using the open-source Biomechanical Toolkit library Raw EMG signals were band-pass filtered between 10 Hz-450 Hz (Butterworth zero-lag 4^th^ order), full-wave rectified, and low-pass filtered at 10 Hz (Butterworth zero-lag 2^nd^ order). All EMG data were time-normalized to a cycling cycle over 360 points.

During the cycling time-to-task failure test, EMG data were extracted at four time points: 0%, 50%, and 100% isotime, as well as at task failure. For each muscle, condition, and time point, up to 20 consecutive pedaling cycles were initially extracted. Following visual inspection, cycles affected by transient artifacts (e.g., movement-induced spikes or signal disturbances related to cable interference) were excluded (see EMG data curation). To ensure a fully balanced within-subject design across participants, conditions, muscles, and time points, the minimum common number of artifact-free cycles available across all conditions was determined (16 cycles). Accordingly, exactly 16 complete pedaling cycles were retained for each participant, muscle, condition, and time point for subsequent analyses.

For EMG amplitude normalization, signals were normalized to the maximal value of the mean EMG profile obtained at 0% isotime for each muscle and each condition. Specifically, for each participant, muscle, and condition, the 16 retained pedaling cycles at 0% isotime were averaged to compute a representative mean EMG waveform. The peak amplitude of this mean waveform was then used as the normalization reference for all corresponding time points within the same condition. This procedure ensured within-condition normalization while preserving valid between-condition comparisons and avoiding potential confounding influences associated with maximal voluntary contractions or separate normalization tasks. After amplitude normalization, the full time-normalized EMG waveform of each retained cycle was preserved without temporal averaging across cycles in order to maintain intra-individual and inter-cycle variability. Mean and standard deviation waveforms were computed for descriptive visualization purposes only. The normalized EMG profiles were subsequently analyzed using Statistical Parametric Mapping to compare activation patterns between conditions across the pedaling cycle (see *Statistical Analysis*).

To determine whether mental fatigue influenced the consistency of neuromuscular activation, we performed a variability analysis of the EMG signal. For each muscle, time point (isotime levels and task failure), and condition (Stroop vs. Control), the point-by-point standard deviation of the EMG signal across the 16 retained cycles was calculated over the pedaling cycle. These standard deviation waveforms were used as indices of inter-cycle variability and were interpreted as a measure of the consistency of muscle activation patterns across repeated pedal cycles. Differences in variability between conditions were assessed using SPM (see *Statistical Analysis*). This approach allowed us to examine whether mental fatigue altered the stability and temporal consistency of muscle activation across repeated pedal strokes.

### EMG data curation

EMG data were curated prior to statistical analysis to ensure signal quality and balanced statistical comparisons. Electrodes presenting persistent signal loss or severe artifacts in one of the two conditions (e.g., low signal-to-noise ratio, movement-induced interference, or signal dropout during prolonged acquisition) were excluded from analysis in both conditions to maintain within-subject comparability. Across the 340 total muscle recordings (10 muscles × 18 participants × 2 conditions), 21 recordings (6%) were excluded due to poor signal quality. After electrode exclusion, the number of participants retained per muscle for complete 2 conditions was: SOL (n = 17), RF (n = 14), VL (n = 15), BF (n = 13), GL (n = 17), TA (n = 17), VM (n = 17), ST (n = 16), GMax (n = 14), and GM (n = 14). A total of 2720 pedaling cycles were collected across all participants and conditions. Following visual inspection, 83 cycles (3%) were removed due to transient artifacts. After this selection, the minimum common number of valid cycles across all participants, conditions, muscles, and time points was 16 cycles, which were retained for statistical analysis to ensure a fully balanced repeated-measures design.

### Psychological variables

Visual analog scales (VAS) were used to assess participants’ mental fatigue, motivation, and boredom. The VAS was printed as 100-mm horizontal lines, and participants responded to each scale-specific question by placing a vertical mark on the line. For each VAS, the left and right anchors and specific question, as well as the timing of each measurement, are described below.

### Mental fatigue

In the control and Stroop conditions, participants reported their level of mental fatigue before the cognitive tasks, after completing half of the cognitive task (30 min), immediately after the cognitive task, and after the cycling time-to-task failure test. The VAS was anchored with “not fatigued at all” and “extremely fatigued” on the left and right ends of the scale. The following question was positioned above the VAS: *“What is your current level of mental fatigue, which is your feeling of exhaustion and lack of energy?”* (Jacquet et al. 2021).

### Motivation

Because fatigue could influence motivation (Müller and Apps 2019) and that the effort allocated in a task is related to the motivation to perform the task (Richter et al. 2016), we measured changes in motivation. Participants reported their level of motivation related (i) to the cognitive tasks before the tasks, midway through the cognitive tasks (30 min) and immediately after the cognitive tasks; and (ii) the cycling task before and immediately after the task. The VAS was anchored with “not motivated at all” and “extremely motivated” on the left and right ends of the scale. The following question was positioned above the VAS before the cognitive and cycling tasks, halfway through the cognitive tasks, and after the cognitive and cycling tasks, respectively: *“What is your motivation to perform the task at hand?”*, *“What is your motivation to continue the task you have just completed?”*, *“What is your motivation to repeat the task you have just completed?”*

### Boredom

Because boredom has been identified as a confounding factor in the development of mental fatigue (Mangin and Pageaux 2025), we measured changes in boredom during the cognitive tasks. Participants reported their level of boredom midway through the cognitive tasks and immediately after the cognitive tasks. The VAS was anchored with “not bored at all” and “extremely bored” on the left and right ends of the scale. The following question was positioned above the VAS: *“Were you bored during the task you just completed?”*

During the cycling warm-up and time to task failure tests, participants reported the intensity of their perceived effort and muscle pain using the CR100 scale (Borg and Kaijser 2006) at specific and consistent time points: 110 s after the start of the warm-up at 40% PPO, 30 s after starting the time to task failure test at 80% PPO, and every 2 min afterwards and at task failure. Details for each perception are presented below.

### Perceptions of effort

Participants rated the intensity of their perceived effort to cycle. They were asked to refer to their sensations of how hard and intense it felt to move their legs and breathe. Participants were explicitly instructed to consider only their perception of effort and not any other exercise-related perceptions such as muscle pain or fatigue (Bergevin et al. 2023; Pageaux 2016). The CR100 scale ranges from 0 (no effort at all) to 100 (maximal effort), with maximal effort anchored to the most intense exertion the participant had ever experienced during a physical task. Modifications were made to the scale’s 0 and 100 anchors to explicitly incorporate the term “effort” (Jacquet et al. 2021). The scale includes verbal anchors corresponding to different intensity levels, such as weak/light (score of 13), moderate (23), strong/heavy (50), and very strong (70). Participants could report values above 100 if they perceived an effort exceeded their maximal effort previously experienced (i.e., rating 100). To enhance understanding, each participant was provided with verbal examples of “no effort at all” (e.g., sitting and relaxing without cognitive engagement) and “maximal effort” (e.g., the most intense effort they had ever experienced during a high intensity whole-body physical task).

### Perceptions of muscle pain

As muscle pain is known to influence endurance performance and to develop during endurance exercise (O’Connor and Cook 1999), we assessed muscle pain during the cycling task for exploratory purposes. Participants used a separate modified CR100 scale to assess muscle pain, with the word “pain” explicitly incorporated into the anchor descriptions: 0 (no pain at all) and 100 (maximal pain). Similar to the perceived effort scale, memory anchoring was used to define the 100-point reference. Participants were instructed to rate “the intensity of pain they felt during the exercise”, focusing specifically on localized sensations in the leg muscles. These sensations were described as potentially including feelings of burning, intense, or sharp pain (O’Connor and Cook 1999). It was emphasized that participants should not confuse or equate their ratings of effort with their perception of pain (Bergevin et al. 2023; Pageaux 2016). To ensure proper use of the scale, each participant was provided with verbal examples illustrating “no muscle pain” and “maximal muscle pain” as reference points.

### Subjective workload

Participants reported their perceived workload at the end of each experimental condition using the NASA-TLX scale (Hart and Staveland 1988), that consists of six items, i.e., (i) mental demand, (ii) physical demand, (iii) temporal demand, (iv) performance, (v) effort, and (vi) frustration. Participants rated each item on a scale divided into twenty equal intervals anchored by a bipolar descriptor (i.e., high/low). This score was multiplied by five, resulting in a final score between 0 and 100 for each item.

### Statistics

All data are presented as means ± standard deviation, unless stated otherwise. The normality of the data was verified using the Shapiro-Wilk test and visual inspection of QQ plots. Statistical significance was set at α ≤ 0.05 for all analyses. Effect sizes are reported as partial eta squared (*η_p_^2^*) for ANOVAs, with thresholds for small, moderate, and large effects set at 0.01, 0.07, and 0.14, respectively (Cohen 1988). Cohen’s d was calculated for t-tests, with thresholds for small, moderate, and large effects set at 0.2, 0.5, and 0.8, respectively (Cohen 1988). When a Wilcoxon signed-rank test was performed, the rank biserial correlation (r_rb) is reported. Post-hoc comparisons from repeated measures ANOVAs were corrected for multiple comparisons using the Holm-Bonferroni method. Greenhouse-Geisser correction was applied to degrees of freedom when violations of sphericity were detected. Except for EMG data analyzed with Matlab, all statistical analyses were conducted using Jamovi 1.2.27.

### Time to task failure

Paired *t*-tests were conducted to assess differences in endurance performance between conditions (Stroop vs. Control). Where data deviated from normality, a Wilcoxon signed-rank test was additionally performed, and results of both tests are reported.

#### EMG

To assess differences in muscle activation patterns between Stroop and Control conditions, we performed time-series analyses using Statistical Parametric Mapping (SPM1D v0.4 Matlab library) (Pataky et al. 2013). EMG signals and EMG signals standard deviations were analyzed across the normalized pedaling cycle (0–100%), with separate analyses conducted for each muscle. For the isotime series (0%, 50%, and 100%) during the time to task failure test, a two-way repeated-measures ANOVA was used, with condition (Stroop vs. Control) and isotime point (three isotime levels) as within-subject factors. This analysis allowed us to evaluate the evolution of muscle activation across the task duration and whether this evolution differed between Stroop and Control conditions under increasing fatigue. In addition, paired t-tests were conducted using SPM1D to specifically compare both conditions at task failure. Isolating these comparisons enabled targeted hypothesis testing without unnecessarily complicating the factorial design. Each analysis was based on 16 complete pedaling cycles per subject, per condition, and per time point, with no temporal averaging, to preserve within-subject variability and cycle-to-cycle dynamics. Muscles were analyzed individually, rather than as a factor in the statistical models, based on physiological and methodological rationale. Each muscle has a distinct activation profile and biomechanical function during cycling, and comparing signals with fundamentally different temporal shapes would violate the assumptions of time-series based SPM analysis. Treating each muscle independently ensured both physiologically meaningful and statistically appropriate comparisons. Statistical significance was set at α = 0.05 for all SPM analyses.

### Perceptual responses

Paired t-tests were conducted to assess the effect of condition on perceptions of effort and muscle pain recorded during the warm-up and at task failure. A two-way repeated measures ANOVA [condition (Stroop vs. Control) × time (0%, 50%, and 100% isotime)] was conducted for perceptual responses recorded during the time to task failure test. Where data deviated from normality, Wilcoxon signed-rank tests were additionally performed.

### Mental fatigue, motivation and boredom

Repeated measures ANOVAs were used to assess changes over time in these variables within each condition. For mental fatigue, the factor time included four levels: pre cognitive task, midway through the cognitive task, post cognitive task, and post cycling. For motivation to perform the cognitive tasks, the factor time included three levels: pre cognitive task, midway through the cognitive task, and post cognitive task. For motivation to perform the cycling tests, the factor time included two levels: pre cycling and post cycling. For boredom, the factor time included two levels: midway through the cognitive task and post cognitive task.

## Results

### Manipulation checks for mental fatigue induction

*Mental fatigue*. Mental fatigue increased overtime (*F*(2, 34) = 29.19, *p* < .001, *n_p_^2^* = .632) and was higher in the Stroop than in the Control condition (main effect of condition: *F*(1, 17) = 8.36, *p* = .010, *n_p_^2^* = .330). The condition × time interaction reached significance (*F*(3, 51) = 4.02, *p* = .012, *n_p_^2^* = .191), and the details of the posthoc comparisons are presented in Figure 2A. Participants reported similar levels of mental fatigue pre cognitive task in the Stroop and Control condition (*t*(17) = -0.879, *p* = .783, *d* = .207, *95%CI* [-.671, .263]), but higher levels of mental fatigue post cognitive task in the Stroop compared to the Control condition (*t*(17) = -3.55, *p* = .015, *d* = -.837, *95%CI* [-1.367, -.288]).

**Figure 2.**
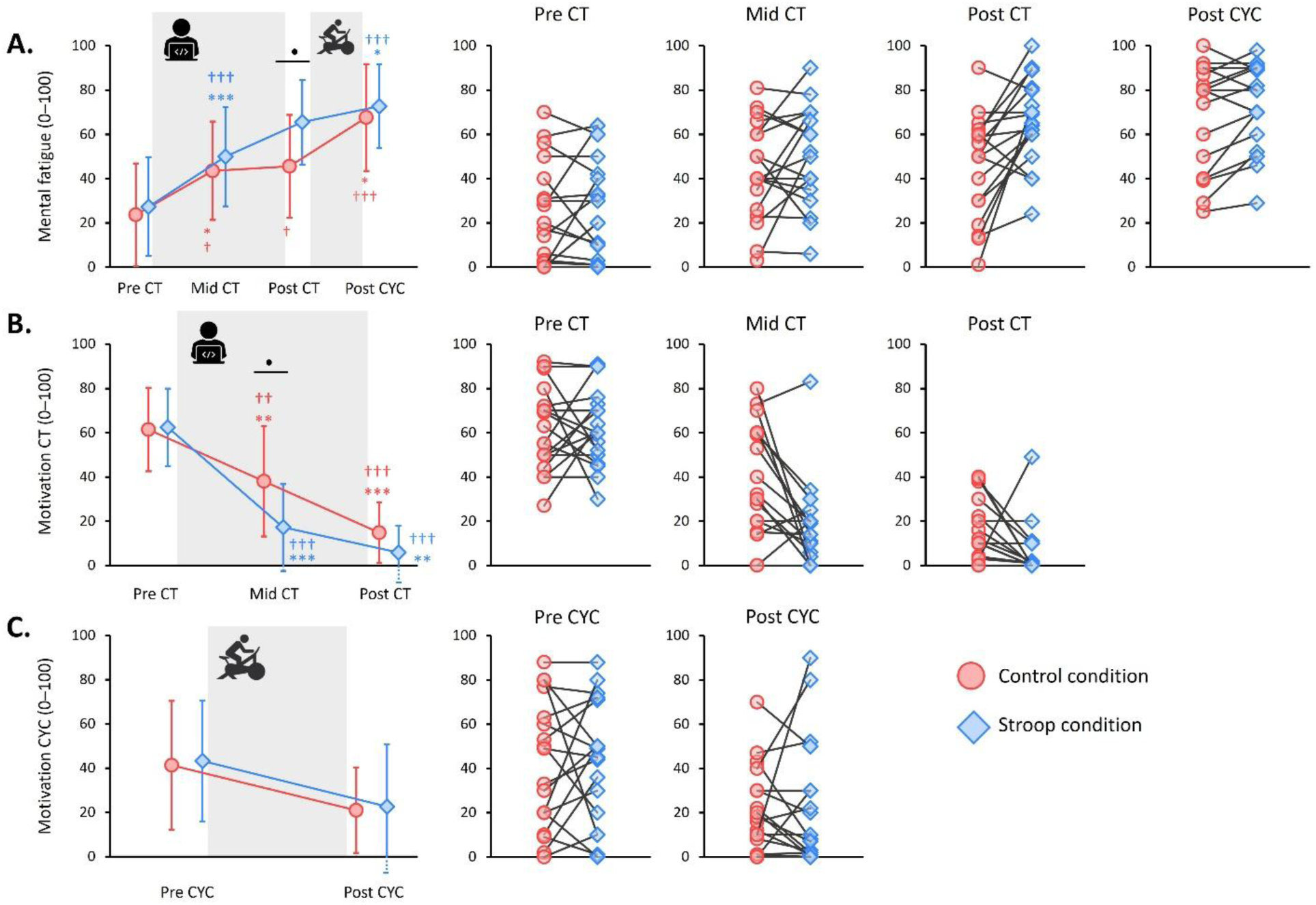
Changes in mental fatigue and motivation during the control (documentary) and Stroop conditions. Panel A shows mean and individual values for mental fatigue level before the cognitive tasks (pre CT), midway through the cognitive tasks (mid CT), immediately after the cognitive tasks (post CT) and after the cycling time to task failure test (post CYC). Panel B shows mean and individual values for motivation to perform the cognitive task measured pre CT, mid CT and post CT. Panel C shows mean and individual values for motivation to perform the cycling time to task failure test measured pre CYC and post CYC. The symbol • indicates a difference between conditions at the same time point; * indicates a difference with the previous time point within the same condition; † indicates a difference with pre CT within the same condition. One symbol denotes *p* < .05, two symbols denote *p* < .01, three symbols denote *p* < .001. Data are presented as mean ± standard deviation.

### Motivation

Motivation to perform the cognitive tasks decreased overtime (*F*(2, 34) = 94.36, *p* < .001, *n_p_^2^* = .847) and was lower in the Stroop than in the Control condition (main effect of condition: *F*(1, 17) = 5.88, *p* = .027, *n_p_^2^* = .257). The condition × time interaction reached significance (*F*(2, 34) = 5.99, *p* = .006, *n_p_^2^* = .261), and details of the posthoc results are presented in Figure 2B. Pre cognitive task, participants reported similar levels of motivation in the Stroop and Control conditions (*t*(17) = -.175, *p* = .863, *d* = -.041, *95%CI* [-.503, .422]). Midway through cognitive tasks, participants were more motivated to continue in the Control than in the Stroop condition (*t*(17) = 3.382, *p* = .011, *d* = .797, *95%CI* [.256, 1.321]). Post cognitive tasks, participants reported similar levels of motivation to repeat the cognitive task they had just completed (*t*(17) = 2.027, *p* = .117, *d* = .478, *95%CI* [-.017, .960]). Motivation (Figure 2C) to perform the cycling tests decreased from pre to post test (*F*(1, 17) = 12.49, *p* = .003, *n_p_^2^* = .424), with no difference between conditions (*F*(1, 17) = .108, *p* = .476, *n_p_^2^* = .006) and no condition × time interaction (*F*(1, 17) = .003, *p* = .956, *n_p_^2^* < .001).

### Boredom

VAS scores for boredom were 4.0 ± 2.8 and 4.3 ± 3.0 at midway and post task for the Control condition, and 5.3 ± 2.5 and 6.5 ± 2.7 for the Stroop condition. Participants were more bored when performing the Stroop task than when watching the documentary (main effect of condition: *F*(1, 17) = 6.61, *p* = .020, *n_p_^2^* = .280). Boredom increased from midway to post task (*F*(1, 17) = 7.80, *p* = .012, *n_p_^2^* = .314), with no condition × time interaction (*F*(1, 17) = 2.39, *p* = .140, *n_p_^2^* = .123).

### Cycling time to task failure

There was no order effect, with the time to task failure in the first condition (793 ± 114 s) not differing from the time to task failure in the second condition (812 ± 199 s) (*t*(17) = 0.36, *p* = .726, *d* = -.121, *95%CI* [-.773, .539]).

The time to task failure decreased by 9.3 ± 11% (−111 ± 160 s) in the Stroop (887 ± 284 s) compared with the Control condition (999 ± 379 s) (t(17) = 2.95, p = .009, d = .694, 95% CI [.169, 1.200]). As the data deviated from a normal distribution, a Wilcoxon signed-rank test was additionally performed as a confirmatory analysis and yielded consistent results (W = 159, p = .001, r_rb = .860). In 83% of participants (15 out of 18), endurance performance was shorter in the Stroop compared with the documentary condition (Figure 3). One participant was identified as a statistical outlier based on the ±2.5 SD criterion applied to the difference scores (Control: 2139 s; Stroop: 1492 s; difference = 647 s, −30%). %). Excluding this participant did not alter the results, with the difference between sessions remaining statistically significant and associated with a large effect size (t(16) = 2.95, p = .002, d = .879, 95% CI [.306, 1.430]).

**Figure 3.**
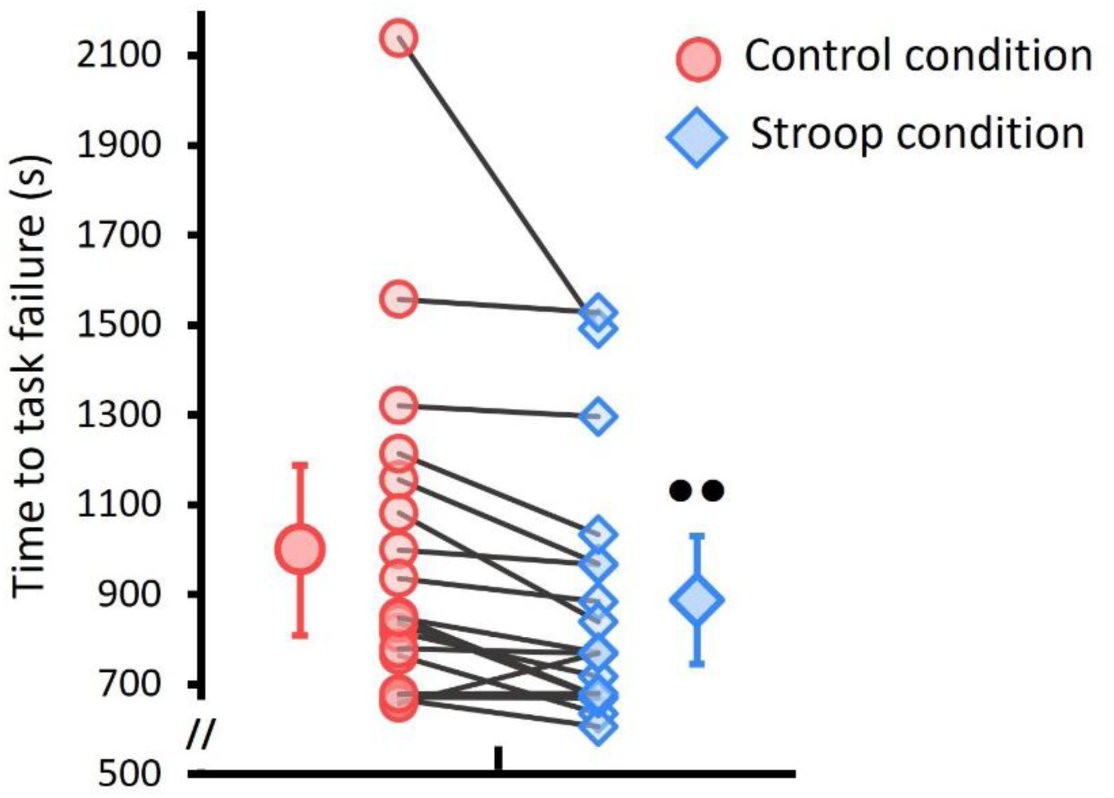
Cycling endurance performance. The cycling time to task failure at 80% peak power output is displayed for the Control (documentary) and Stroop conditions. Large circles and diamonds represent mean data ± standard deviation, while small circles and diamonds indicate individual data points connected by solid lines. •• denotes a significant difference between conditions (*p* < 0.01).

### Electromyography

SPM analyses revealed no significant main effect of condition (Stroop vs. Control) on EMG activation profiles for any muscle at any of the examined time points (Figure 4 and Supplementary Material S1). This absence of difference was also observed for both the time-specific comparison at task failure, with no significant condition-related detected for any muscle (SPM statistic = t; df = 16; *p* > .05; clusters = none) (Figure 5). Similarly, in the two-way repeated-measures ANOVA conducted across the isotime points (0%, 50%, and 100%), the main effect of condition remained non-significant, and no significant condition × isotime point interaction was observed (Table 1). In contrast, a significant main effect of isotime point was observed in specific phases of the pedaling cycle in a subset of muscles: GMax, VL, ST, and GM (Figure 4, Table 1). For clarity reason, data for the RF, BF, VM, SOL and GL are presented as supplementary material S1. Individual data for EMG analysis are provided as supplementary material S2.

**Figure 4.**
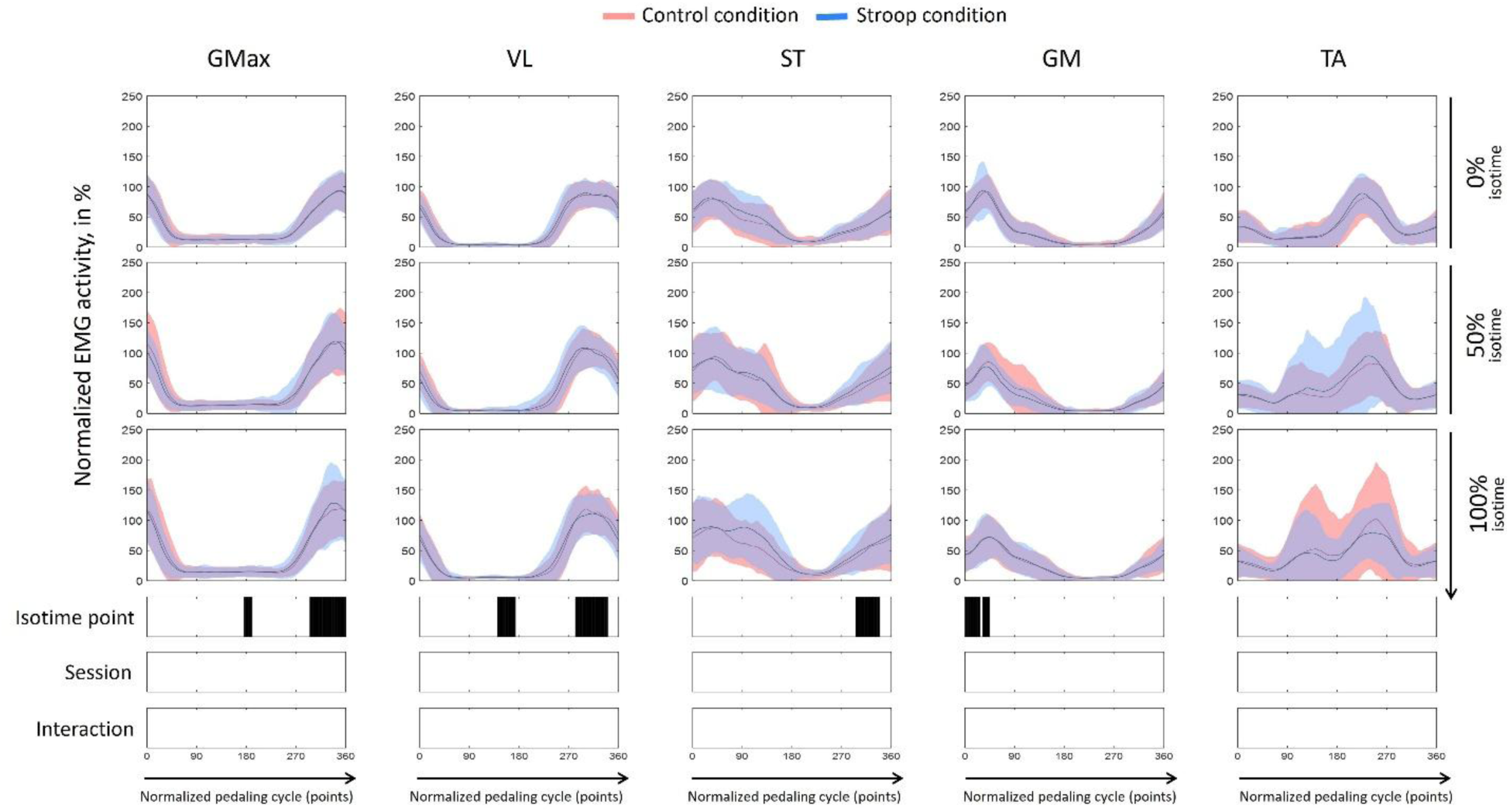
Electromyographic (EMG) activity of the lower limb muscles during the cycling time to task failure test for the control (documentary) and Stroop conditions. Electromyographic (EMG) activity of the gluteus maximus (GMax), vastus lateralis (VL), semi tendinous (ST), gastrocnemius medialis (GM) and tibialis anterior (TA) muscles throughout the pedaling cycle. EMG activity was measured at three isotime points (0%, 50%, and 100% – from top to bottom). Data are presented for two conditions: control (solid red line) and Stroop (solid blue line). The x-axis represents the normalized pedaling cycle (points), while the y-axis indicates normalized EMG activity. Solid lines represent the average muscle activity, with shaded areas denoting the standard deviation. Significant differences (*p* < 0.05) over time, assessed using a two-way repeated-measures ANOVA with factors task condition (Stroop vs. Control) and isotime (0%, 50%, and 100%), combined with Statistical Parametric Mapping analysis, are indicated by vertical black bars beneath each graph.

**Figure 5.**
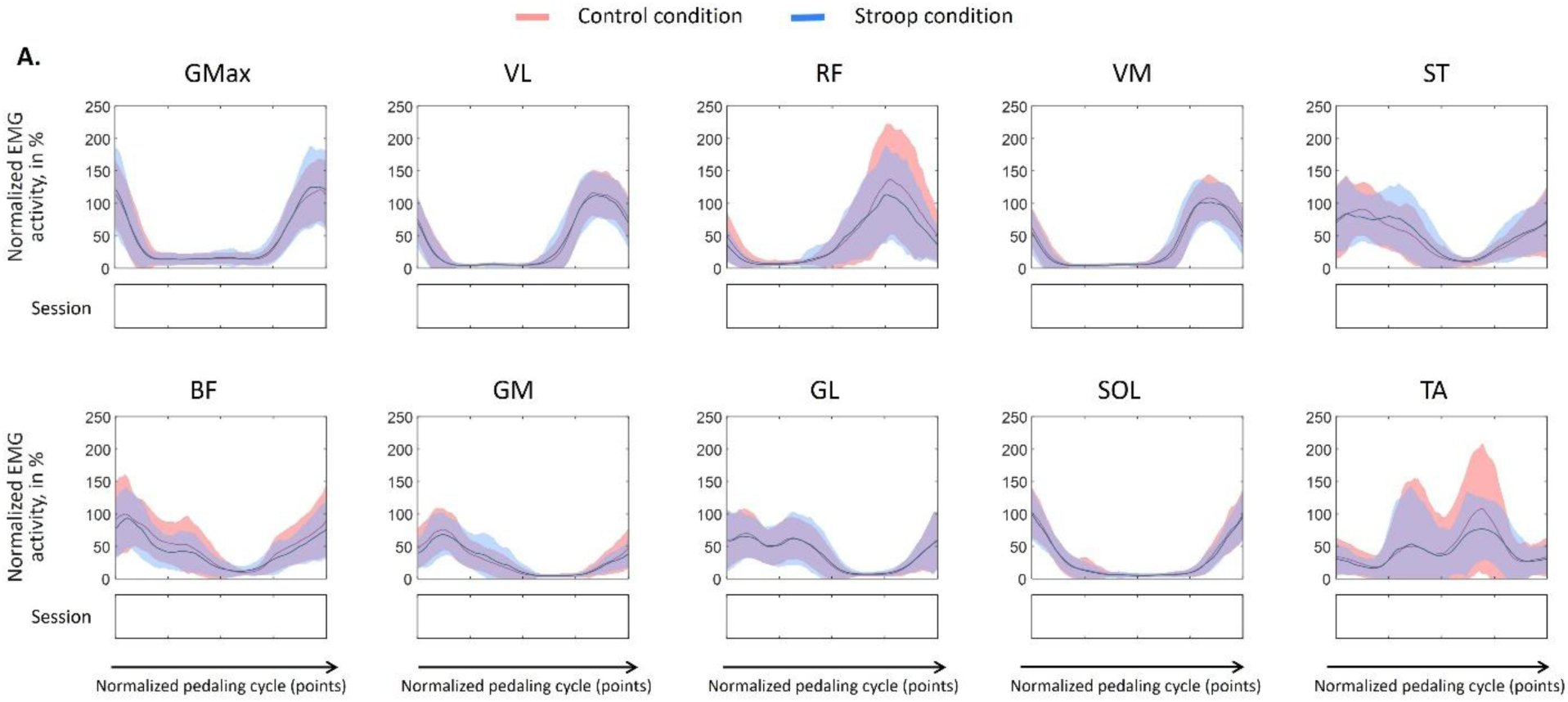
Electromyographic (EMG) activity of the lower limb muscles at task failure for the control (documentary) and Stroop conditions. EMG activity of the gluteus maximus (GMax), vastus lateralis (VL), rectus femoris (RF), vastus medialis (VM), semi tendinous (ST), biceps femoris (BF), gastrocnemius medialis (GM), gastrocnemius lateralis (GL), soleus (SOL) and tibialis anterior (TA) muscles throughout the pedaling cycle. Data are presented for two conditions: control (solid red line) and Stroop (solid blue line). The x-axis represents the normalized pedaling cycle (points), while the y-axis indicates normalized EMG activity. Solid lines represent the average muscle activity, with shaded areas denoting the standard deviation. Statistical differences were calculated using a paired t-test with task condition (Stroop vs. control) combined with Statistical Parametric Mapping analysis.

**Table 1.**
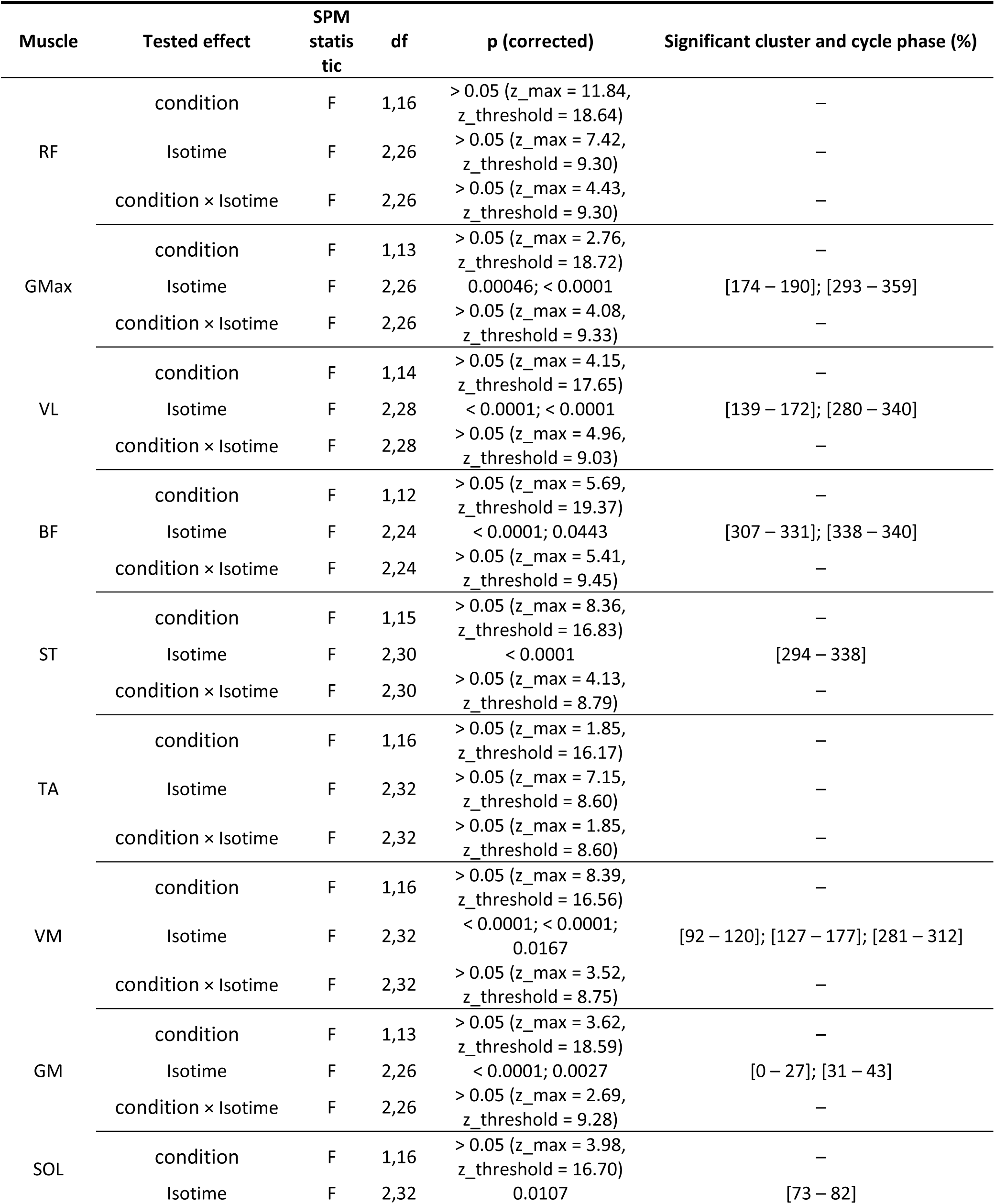
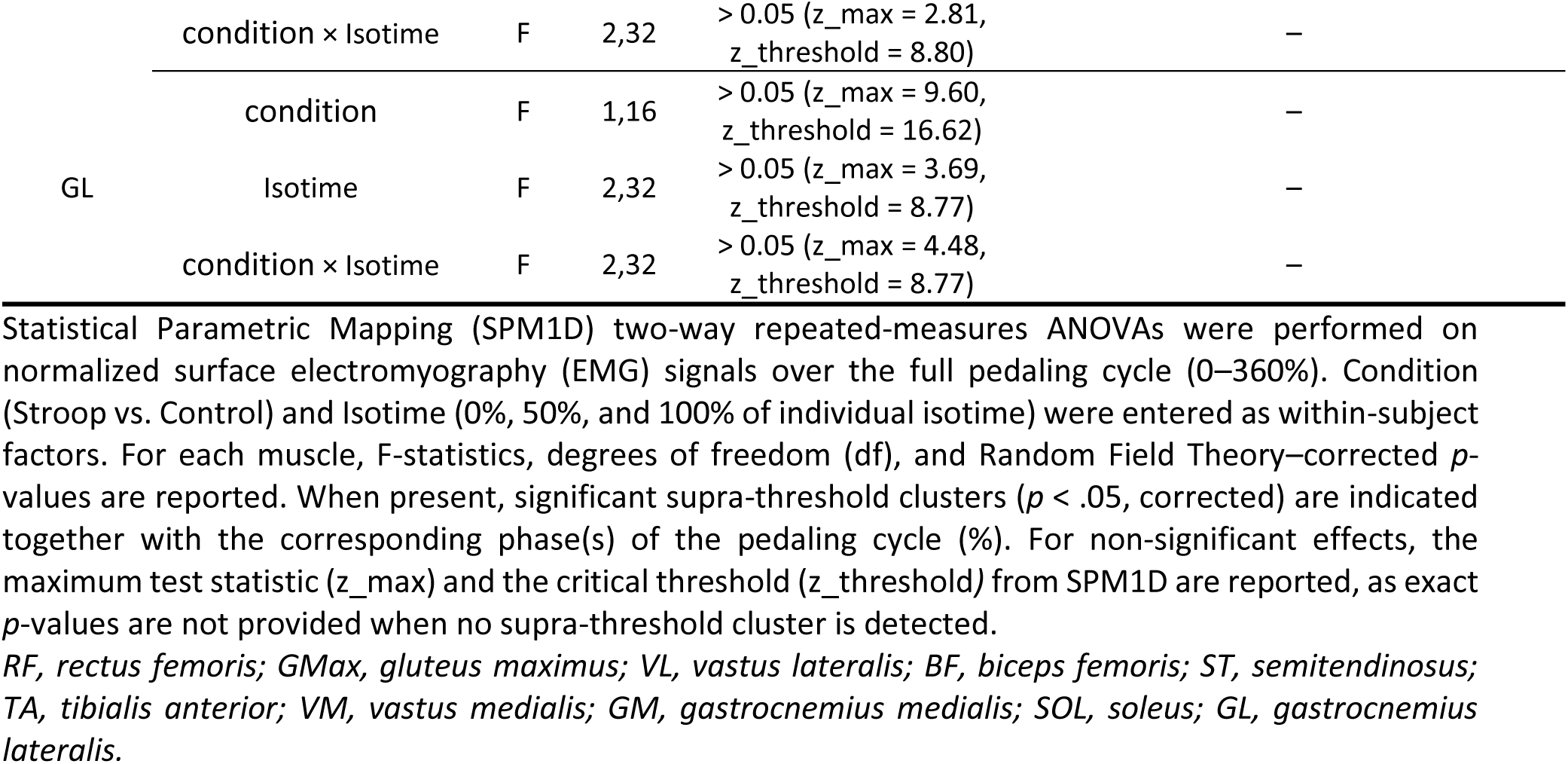
SPM1D results for EMG activity during the cycling time-to-task failure test (isotime analysis)

The variability analysis conducted on the standard deviation profiles of the EMG signals revealed no differences in inter-cycle EMG variability between the Stroop and Control conditions, for any muscle and at any of the tested time points (task failure, and the three isotime points) (Supplementary Materials S3: SPM1D results, S4: isotime data, and S5: task failure data). These results suggest that mental fatigue did not affect the consistency or temporal stability of muscle activation patterns during the cycling test, providing no evidence of altered neuromuscular control stability under mental fatigue.

### Perceptual responses

*Perception of effort.* Mean and individual data are presented in Figure 6A. Participants perceived the warm-up more effortful in the Stroop compared to the Control condition (*t*(17) = -3.149, *p* = .006, *d* = -.742, *95%CI* [-1.258, -.210]). Due to deviation from a normal distribution, we also employed a Wilcoxon rank test which confirmed the difference between condition (*W* = 4.50, *p* = .003, *r_rb_* = -.915). During the time to task failure test, analysis of the isotime values revealed a higher perception of effort in the Stroop compared to the Control condition (main effect of condition: *F*(1, 17) = 7.18, *p* = .016, *n_p_^2^* = .297). The perception of effort increased overtime (*F*(2, 34) = 133.09, *p* < .0.001, *n_p_^2^* = .887), with no condition × time interaction (*F*(2, 34) = 1.00, *p* = .377, *n_p_^2^* = 0.056). At task failure, participants reported the same intensity of perceived effort between conditions (*t*(17) = -1.016, *p* = .324, *d* = -.239, *95%CI* [-.705, .233]) .

**Figure 6.**
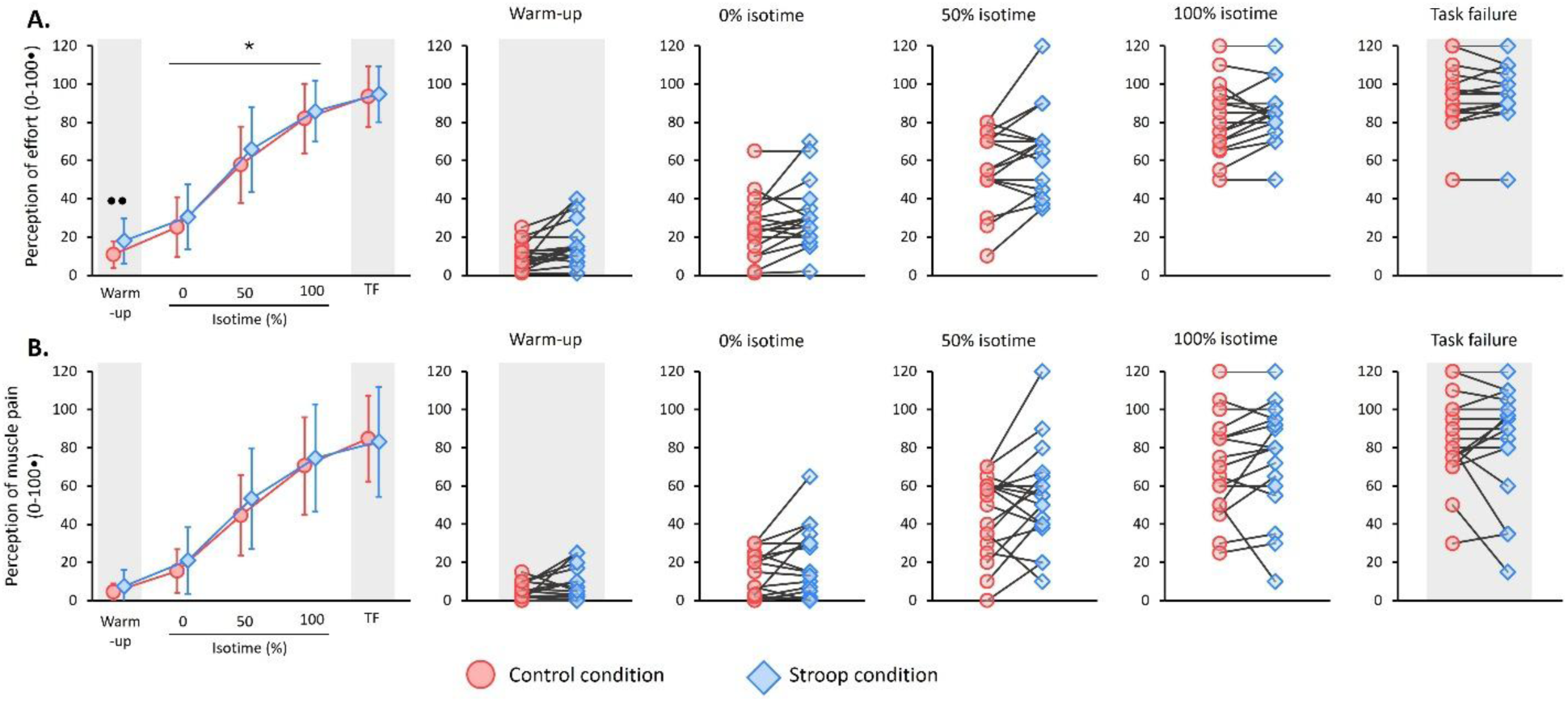
Perceptions of effort and muscle pain during the cycling exercise for the control (documentary) and Stroop conditions. Panel A shows mean and individual values for the perception of effort during the warm-up performed at 40% peak power output (PPO) and the time to task failure test at 80% PPO. During the time to task failure test, data are presented at four time points: 0%, 50% and 100% individual isotime, and at task failure. Panel B shows mean and individual values for the perception of muscle pain at the same time points. For clarity, the warm-up and task failure phases fall into a grey area: the warm-up was performed at a different pedaling intensity (40% PPO), and the task failure data were collected at a non-isotime time point. The symbol • indicates a difference between conditions at the same time point; * indicates a main effect of condition during the isotime. One symbol denotes *p* < .05, two symbols denote *p* < .01. Data are presented as mean ± standard deviation.

### Perceptions of muscle pain

Mean and individual data are presented in Figure 6B. During the warm-up, participants reported a similar level of muscle pain between the Stroop and Control conditions (*t*(17) = -1.550, *p* = .140, *d* = -.365, *95%CI* [-.838, .118]). Due to deviation from a normal distribution, we also employed a Wilcoxon rank test, which also suggested no difference between conditions (*W* = 35.50, *p* = .299, *r_rb_* = -.324). During the time to task failure test, analysis of the isotime values suggested a higher muscle pain intensity in the Stroop compared to the Control condition, without reaching significance (main effect of condition: *F*(1, 17) = 3.952, *p* = .063, *n_p_^2^* = .189). Muscle pain increased overtime (*F*(2, 34) = 90.023, *p* < .001, *n_p_^2^* = .841), with no condition × time interaction (*F*(2, 34) = .693, *p* = .507, *n_p_^2^* = .039). At task failure, participants reported the same intensity of muscle between conditions (*t*(17) = .373, *p* = .713, *d* = .088, *95%CI* [-0.376, 0.550]). Due to deviation from a normal distribution, we also employed a Wilcoxon rank test, which also suggested no difference between conditions (*W* = 34.00, *p* = .964, *r_rb_* = -.030).

#### NASA-TLX

The Control and Stroop conditions differed on the following NASA-TLX items: mental demand (58 ± 32 vs. 81 ± 16, respectively; *p* = .007, *d* = .73, *95%CI* [.198, 1.240]), performance (29 ± 24 vs. 39 ± 27; *p* = .049, *d* = .49, *95%CI* [-.008, .983]), and frustration (34 ± 26 vs. 49 ± 32, respectively; *p* = .008, *d* = .70, *95%CI* [.176, 1.210]). No difference was reported for the other items related to physical and temporal demand, and effort (all *p* > .05, *d* = .14–.38).

## Discussion

This study examined the effects of mental fatigue induced by a prolonged demanding cognitive task, compared with a control task, on cycling endurance performance, effort perception and surface EMG-derived muscle activation. Mental fatigue reduced endurance performance and increased the perception of effort, but did not alter EMG-related parameters during the cycling endurance task.

### No EMG Evidence of Altered Motor Command Under Mental Fatigue

Our first objective was to investigate whether the increased perceived effort in the presence of mental fatigue was associated with alterations in muscle activation during cycling. Our hypothesis of altered muscle activation was grounded on prior evidence reporting changes in EMG activity following prolonged mental exertion during upper- and lower-limb isometric tasks (Bray et al. 2008; Graham et al. 2014; Jacquet et al. 2021; Budini et al. 2014), as well as during a submaximal cycling task (Pageaux et al. 2015). Collectively, these findings suggest that mental fatigue may affect central motor command during subsequent physical tasks, potentially contributing to the observed increased perceived effort.

Because cycling requires the coordinated activation of multiple muscles (So et al. 2005), the study by Pageaux et al. (2015), which focused on a single muscle, does not provide a comprehensive understanding of the effects of mental fatigue on muscle activation during a whole-body endurance task. Therefore, we aimed to extend these findings by examining muscle activation using surface EMG recordings from ten lower-limb muscles to better capture peripheral indices of motor command. Contrary to our hypothesis, mental fatigue did not alter muscle activation patterns. This result aligns with a recent work that reported no changes in high density-EMG-derived neurophysiological outcomes (i.e., motor unit activity, common synaptic inputs) following a 30-min Stroop task and a subsequent isometric contraction to failure (Alix-Fages et al. 2023). Given that our study was powered to detect medium-to-large effect, our results, together with those of Alix-Pages et al. (2023), suggest that mental fatigue either does not affect EMG-derived measures of muscle activation, or that its effects are small.

Several interpretations can be proposed to account for the absence of significant effects on EMG parameters. First, neural control differs between single-joint and whole-body exercises (Sidhu et al. 2013a). The dynamic nature of cycling likely leads to EMG patterns distinct from those observed during isometric, single-joint tasks and may permit compensatory strategies that are not detectable through mean EMG amplitude alone (Bray et al. 2008; Graham et al. 2014; Jacquet et al. 2021). However, we also examined inter-cycle EMG variability as an index of the consistency of muscle activation patterns across repeated pedal cycles. The absence of condition effects on both EMG amplitude and inter-cycle variability suggests that mental fatigue did not measurably alter either the magnitude or the temporal consistency of neuromuscular activation during cycling. Second, EMG-derived measures reflect peripheral manifestations of motor command rather than the initial supraspinal processes involved in motor planning and motor command generation. Because mental fatigue induces changes at the supraspinal level (Tanaka et al. 2012; Ishii et al. 2014; Brownsberger et al. 2013; Wiehler et al. 2022; Marcora et al. 2009; Pageaux et al. 2013), it is possible that compensatory adjustments occur during the early stages of motor command planning or generation, before transmission through the spinal cord. In such a case, muscle activation could remain unchanged despite altered supraspinal activity. Accordingly, while our results suggest that the neural information reaching the muscles is similar in the presence and absence of mental fatigue, alterations at earlier stages of motor command generation cannot be excluded. For instance, it has been reported that continuous repetition of motor imagery inducing mental fatigue decreased corticospinal excitability measured with transcranial magnetic stimulation (Nakashima et al. 2021; Nakashima et al. 2022). Third, the absence of changes in EMG-derived muscle activation suggests that the observed decline in endurance performance is unlikely to primarily result from alterations in motor command. This interpretation aligns with recent evidence suggesting that mental fatigue may instead reflect metabolic alterations within brain regions involved in cognitive control (Pessiglione et al. 2025; Wiehler et al. 2022; Barakat et al. 2025). Because effort is also perceived during cognitive tasks (Mangin and Pageaux 2026) , such metabolic disturbances may increase the perceived cost of effort, thereby contributing to the higher perceived effort observed under mental fatigue. Notably, these metabolic changes have been proposed to increase the cost of cognitive control, biasing decision-making toward low-effort actions associated with immediate rewards (Barakat et al. 2025; Pessiglione et al. 2025). In the context of a cycling time-to-task failure test, such low-effort actions would correspond to premature task disengagement.

### Mental Fatigue Impairs Endurance Performance

Our second objective was to replicate the detrimental effect of mental fatigue on endurance performance. Participants began with similar levels of mental fatigue in both conditions (Control vs. Stroop). However, mental fatigue was significantly higher following 1 h of the Stroop task compared with 1 h of watching a neutral documentary. Because boredom increased to a similar extent in both conditions, the greater rise in perceived mental fatigue observed during the Stroop task cannot be attributed to differences in boredom over time (Mangin and Pageaux 2025), confirming that the Stroop task successfully induced a specific state of mental fatigue compared with the documentary condition.

Interestingly, although motivation to perform the cognitive task decreased to a greater extent in the Stroop condition, participants reported similar levels of motivation to perform the cycling endurance task in both conditions. This observation suggests that the change in motivation observed after the mental fatigue manipulation was task-specific and primarily related to the cognitive task used to induce mental fatigue. This result not only supports the negative effect of mental fatigue on motivation (Müller and Apps 2019), but also suggests that the reduction in motivation induced by mental fatigue may be task-specific and primarily related to the cognitive task used to induce mental fatigue. Furthermore, because participants reported similar levels of motivation to perform the cycling task in each condition, the reduction in endurance performance cannot be attributed to alterations in cycling-specific motivation. Future studies designed to test cost–benefit models of fatigue and motivation could further examine how fatigue, motivation, perceived effort and boredom contribute to the perceived cost of endurance exercise.

Although the effect of mental fatigue on physical performance has recently been questioned (Holgado et al. 2021), we observed, in line with our hypothesis, an impaired cycling endurance performance in the presence of mental fatigue. This finding aligns with previous studies that used the Stroop task to induce mental fatigue and examine its impact on endurance performance (Bray et al. 2008; Smith et al. 2016; Boat et al. 2018; Pageaux et al. 2014). Endurance tasks, such as a time to task failure test as used in the present study, require continuous regulation of effort over time (Pageaux and Lepers 2018; Englert et al. 2021), during which perceived effort plays a central regulatory role (Pageaux 2014). Because mental fatigue is proposed to increase perceived effort, the detrimental effects of mental fatigue are particularly likely to manifest during endurance tasks.

Our results align with this possibility. During the Stroop condition, the elevated perception of effort observed from the onset of the cycling warm-up, along with the significantly higher rate of increase in perceived effort during cycling (Stroop condition : 6.7 ± 2.5, control condition 6.0 ± 1.8 units.min^-1^; p = .031), led to premature task disengagement and decreased endurance performance. These findings reinforce previous evidence on the negative effect of mental fatigue on endurance performance (Marcora et al. 2009; Pageaux 2014; Smith et al. 2016) and provide further experimental that mental fatigue impairs endurance performance primarily through an increased perception of effort.

### Limitations and perspectives for future research

Because our study was powered to detect medium-to-large effects, future studies with larger sample sizes are needed to rule out small effects of mental fatigue on muscle activation assessed using surface EMG. Further, surface EMG provides only a peripheral index of motor command. Given the proposed link between perceived effort and motor command (Bergevin et al. 2023; De Morree et al. 2012), future research should investigate the effects of mental fatigue on corticospinal and intracortical excitability during dynamic exercise using single- and paired-pulse transcranial magnetic stimulation (e.g., Sidhu et al. (2013b)). Assessing corticospinal and intracortical excitability may help determine whether mental fatigue alters processes involved in the transmission of motor command to the working muscles. Furthermore, recent advances in electroencephalography and data analysis have enabled its application during cycling (Pirlot et al. 2025), making it a promising tool for this purpose. In particular, combining electroencephalography with EMG (Petersen et al. 2012) or near-infrared spectroscopy (Zama and Shimada 2015) may help identify supraspinal alterations occurring within prefrontal regions or during early stages of motor command generation in the presence of mental fatigue, or alternatively, provide evidence against their involvement.

Finally, the present study used a deductive approach (Briand et al. 2025) grounded in two theoretical frameworks: one linking perception of effort to endurance performance (Marcora 2019; Marcora et al. 2008; Pageaux 2014), and the other linking perception of effort to motor command (Mangin and Pageaux 2026; Bergevin et al. 2023; de Morree and Marcora 2015). However, we cannot exclude that alternative or complementary mechanisms, grounded in frameworks focusing on interoception, sensory attenuation, or metacognitive inference of fatigue may also contribute to the increased perceived effort and premature task disengagement observed under mental fatigue (Dekerle et al. 2025; Greenhouse-Tucknott et al. 2022; Kuppuswamy 2017). Also, while we did not observe a difference between conditions in motivation to perform the cycling endurance task, we acknlowedge that motivation is a complex phenomenon that may not be fully captured by a single visual analog scale (REF). Therefore, we cannot firmly exclude the possibility that mental fatigue altered effort-based decision-making during endurance exercise through more complex alterations in motivational processes, such as changes in the valuation of the task (Müller and Apps 2019). These frameworks may generate complementary hypotheses regarding sensory information processing sensory information processing, fatigue perception, confidence in the ability to perform the cycling endurance task, and effort-based decision-making during endurance exercise. Because the present study did not include measures specifically designed to assess these processes, our results cannot discriminate between these accounts. Future studies should therefore test these alternative hypotheses using specific a priori designs in the context of endurance exercise and mental fatigue.

## Conclusions

Prolonged cognitive engagement inducing mental fatigue impaired cycling endurance performance by increasing perceived effort. This increased perceived effort occurred in the absence of detectable changes in EMG-derived muscle activation during cycling. This dissociation between perceptual and neuromuscular responses suggests that the detrimental effects of mental fatigue on endurance performance are primarily driven by central mechanisms involved in effort regulation and task disengagement. Future studies should determine whether combining surface EMG with more sensitive neurophysiological approaches can reveal subtle alterations in motor command associated with mental fatigue during dynamic, whole-body exercise.

## CRediT statement

***R. Souron:*** conceptualization; data curation; formal analysis; methodology; supervision; writing- original draft; writing-review & editing.

***A. Sarcher:*** data curation; formal analysis; writing-review & editing.

***L. Lacourpaille:*** conceptualization; methodology; writing-review & editing

***I. Boulahouche:*** data curation; formal analysis; writing-review & editing

***C. Richier:*** data curation; formal analysis; writing-review & editing

***T. Mangin:*** methodology; formal analysis; writing-review & editing

***M. Gruet:*** conceptualization; methodology; writing-review & editing

***J. Doron:*** conceptualization; methodology; writing-review & editing

***M. Jubeau:*** conceptualization; methodology; writing-review & editing

***B. Pageaux:*** conceptualization; methodology; writing-original draft; writing-review & editing

## Use of AI Generated Tools

The authors used ChatGPT (version 5.2, OpenAI) to assist with English language editing and grammar improvement during manuscript preparation. After using this tool, the authors reviewed and edited the content as needed and takes full responsibility for the content of the published article. All scientific content, data interpretation, and conclusions were generated solely by the authors.

