## Supplementary material for "Mental fatigue impairs cycling endurance performance and perception of effort, but not muscle activation": S1

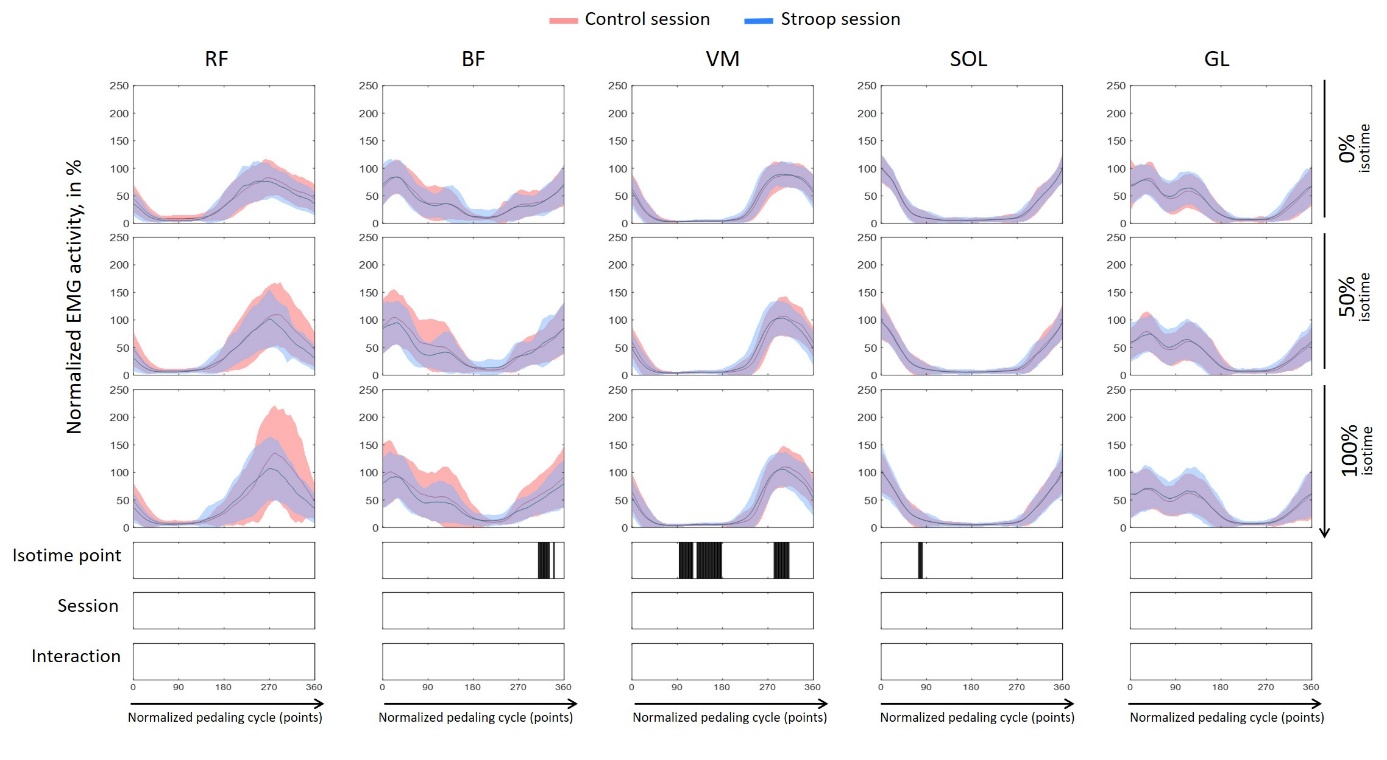

**Supplementary Material S1. Electromyographic (EMG) activity of the lower limb muscles during the cycling time to exhaustion test for the control (documentary) and Stroop sessions**. Electromyographic (EMG) activity of rectus femoris (RF), biceps femoris (BF), vastus medialis (VM), soleus (SOL) and gastrocnemius lateralis (GL) muscles throughout the pedaling cycle. EMG activity was measured at three isotime points (0%, 50%, and 100% – from top to bottom). Data are presented for two sessions: control (solid red line) and Stroop (solid blue line). The x-axis represents the normalized pedaling cycle (points), while the y-axis indicates normalized EMG activity. Solid lines represent the average muscle activity, with shaded areas denoting the standard deviation. Significant differences (*p* < 0.05) over time, assessed using a two-way repeated-measures ANOVA with factors task session (Stroop vs. Control) and isotime (0%, 50%, and 100%), combined with Statistical Parametric Mapping analysis, are indicated by vertical black bars beneath each graph.
