## Supplementary material for "Mental fatigue impairs cycling endurance performance and perception of effort, but not muscle activation": S2

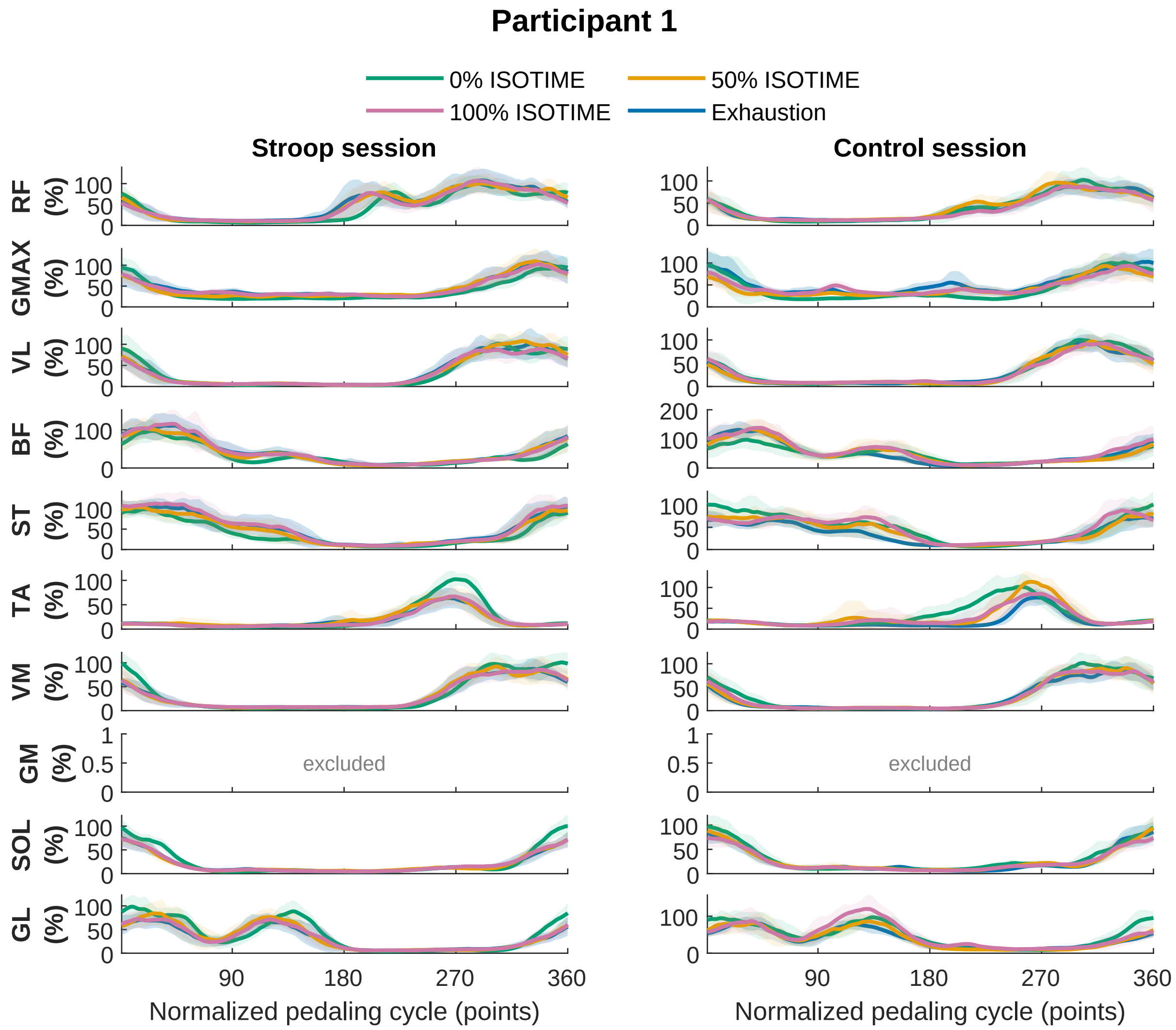
Participant 1
0% ISOTIME
50% ISOTIME
100% ISOTIME
Exhaustion
Stroop session
Control session
RF (%)
GMAX (%)
VL (%)
BF (%)
ST (%)
TA (%)
VM (%)
GM (%)
SOL (%)
GL (%)
excluded
excluded
90
180
270
360
Normalized pedaling cycle (points)
Normalized pedaling cycle (points)

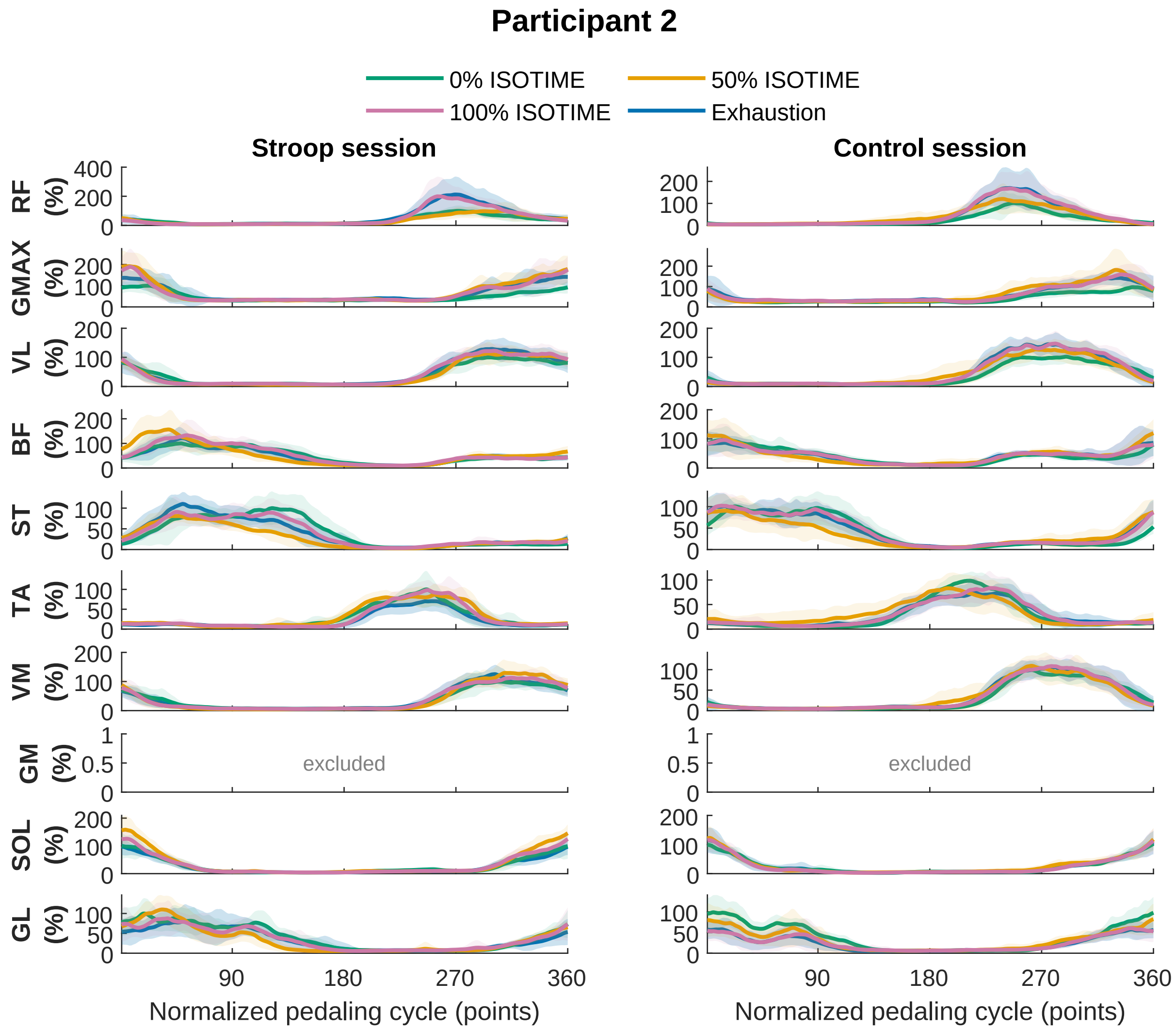

Participant 2
0% ISOTIME
50% ISOTIME
100% ISOTIME
Exhaustion
Stroop session
Control session
RF (%)
GMAX (%)
VL (%)
BF (%)
ST (%)
TA (%)
VM (%)
GM (%)
SOL (%)
GL (%)
excluded
excluded
90
180
270
360
Normalized pedaling cycle (points)
Normalized pedaling cycle (points)

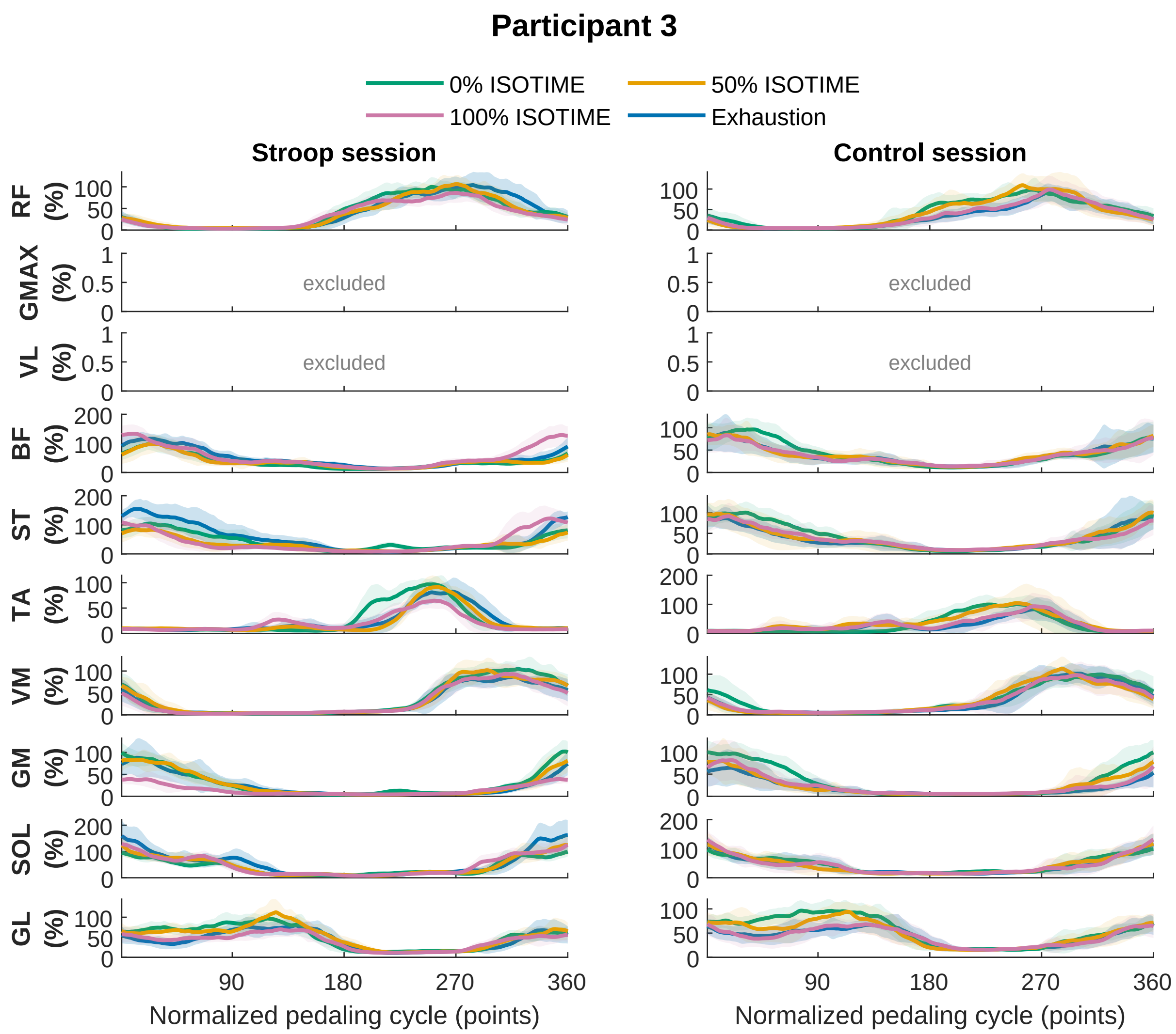

Participant 3
0% ISOTIME
50% ISOTIME
100% ISOTIME
Exhaustion
Stroop session
Control session
RF (%)
GMAX (%)
VL (%)
BF (%)
ST (%)
TA (%)
VM (%)
GM (%)
SOL (%)
GL (%)
excluded
excluded
excluded
excluded
90
180
270
360
Normalized pedaling cycle (points)
Normalized pedaling cycle (points)

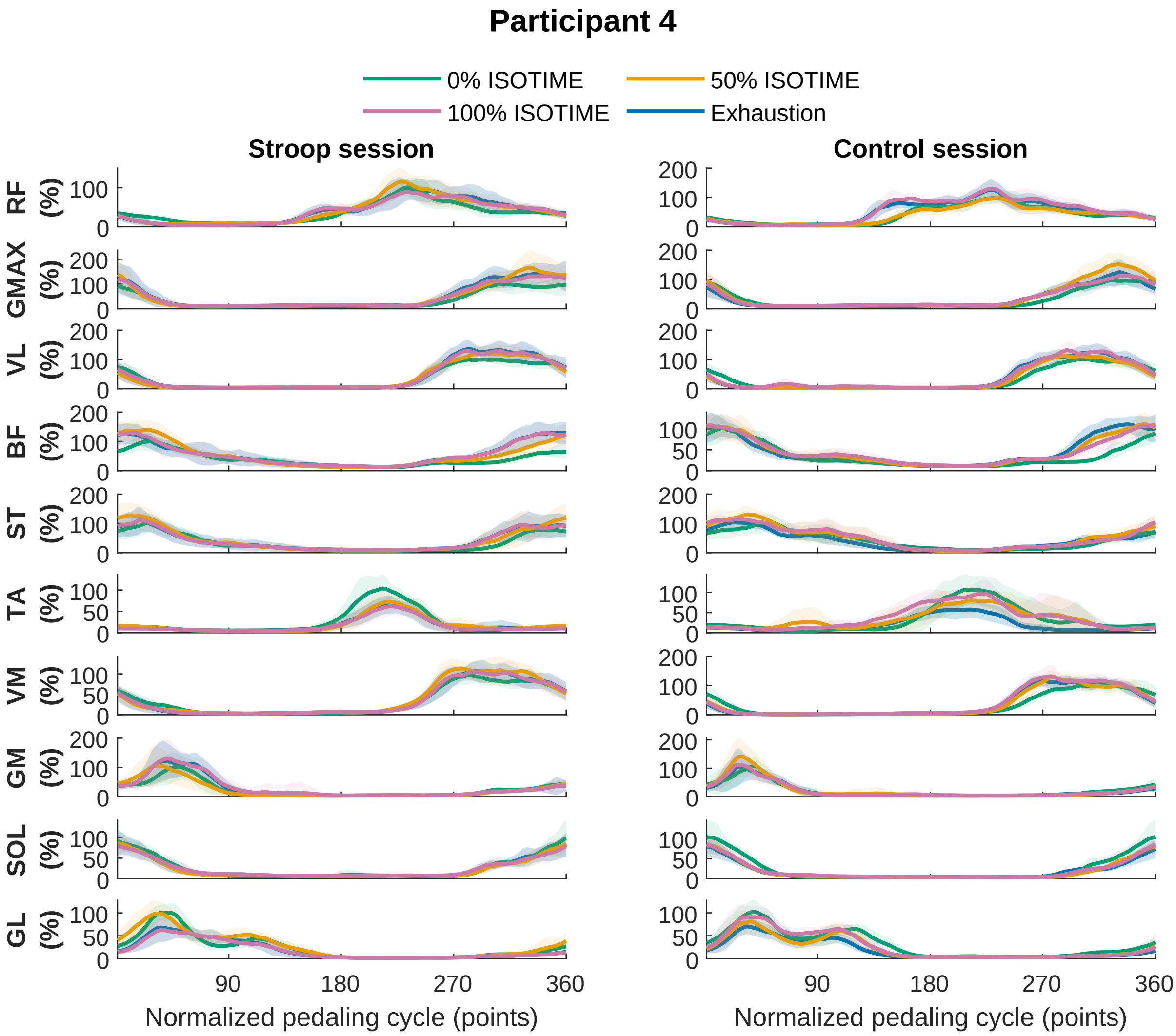

Participant 4
0% ISOTIME
50% ISOTIME
100% ISOTIME
Exhaustion
Stroop session
Control session
RF (%)
GMAX (%)
VL (%)
BF (%)
ST (%)
TA (%)
VM (%)
GM (%)
SOL (%)
GL (%)
90
180
270
360
Normalized pedaling cycle (points)

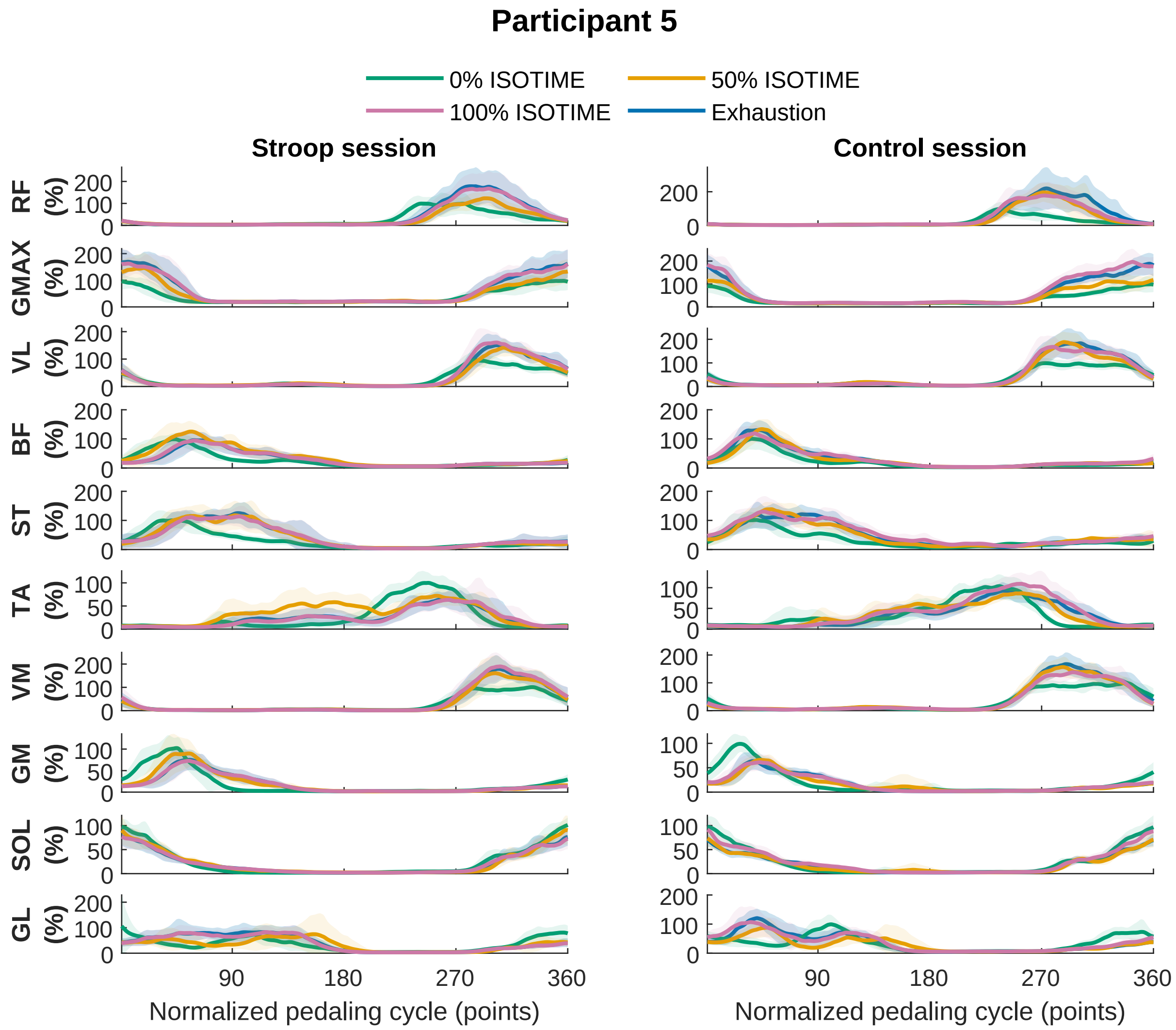

Participant 5
0% ISOTIME
50% ISOTIME
100% ISOTIME
Exhaustion
Stroop session
Control session
RF (%)
GMAX (%)
VL (%)
BF (%)
ST (%)
TA (%)
VM (%)
GM (%)
SOL (%)
GL (%)
90
180
270
360
Normalized pedaling cycle (points)

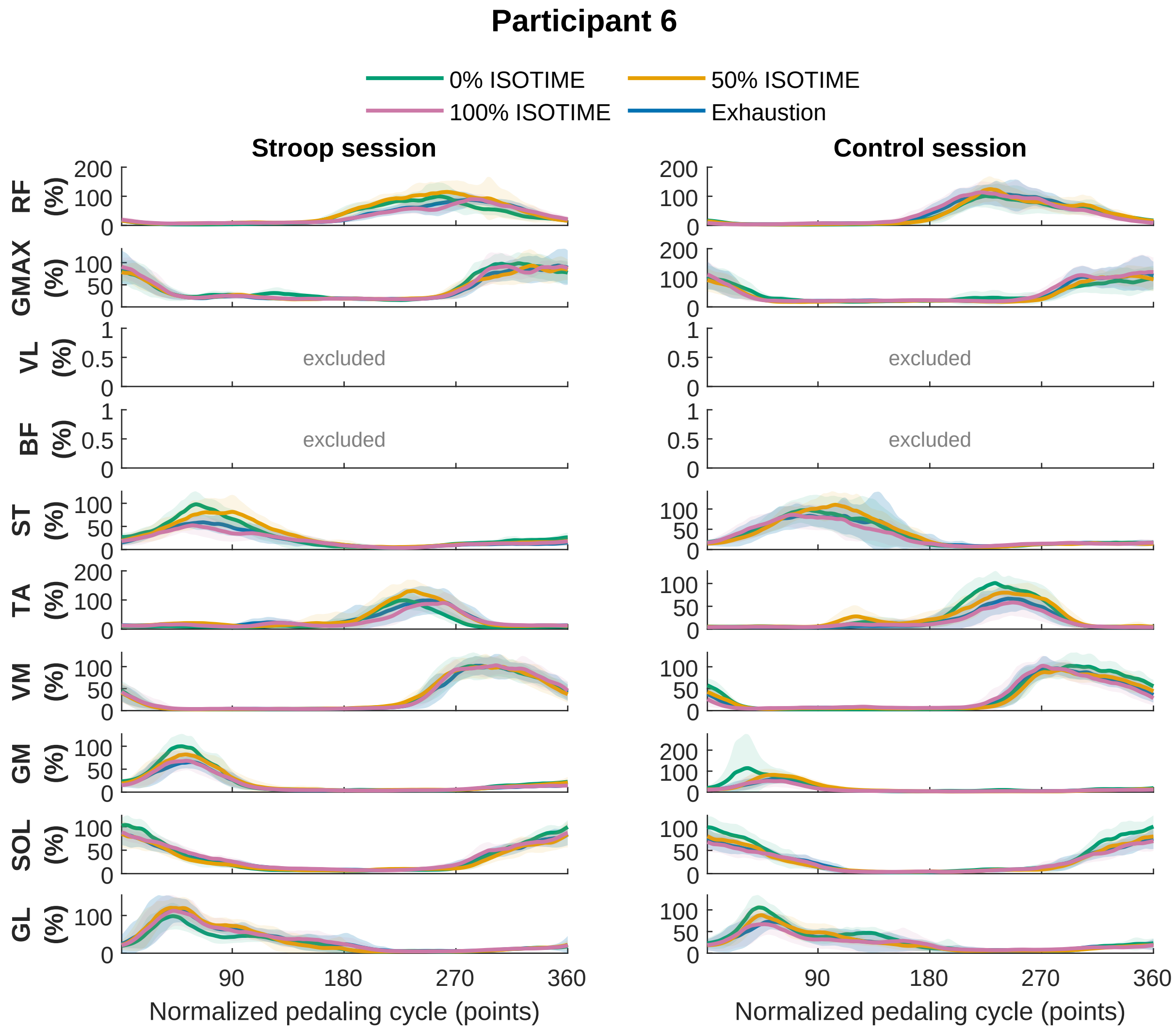

Participant 6
0% ISOTIME
50% ISOTIME
100% ISOTIME
Exhaustion
Stroop session
Control session
RF (%)
GMAX (%)
VL (%)
BF (%)
ST (%)
TA (%)
VM (%)
GM (%)
SOL (%)
GL (%)
excluded
excluded
excluded
excluded
90
180
270
360
Normalized pedaling cycle (points)
Normalized pedaling cycle (points)

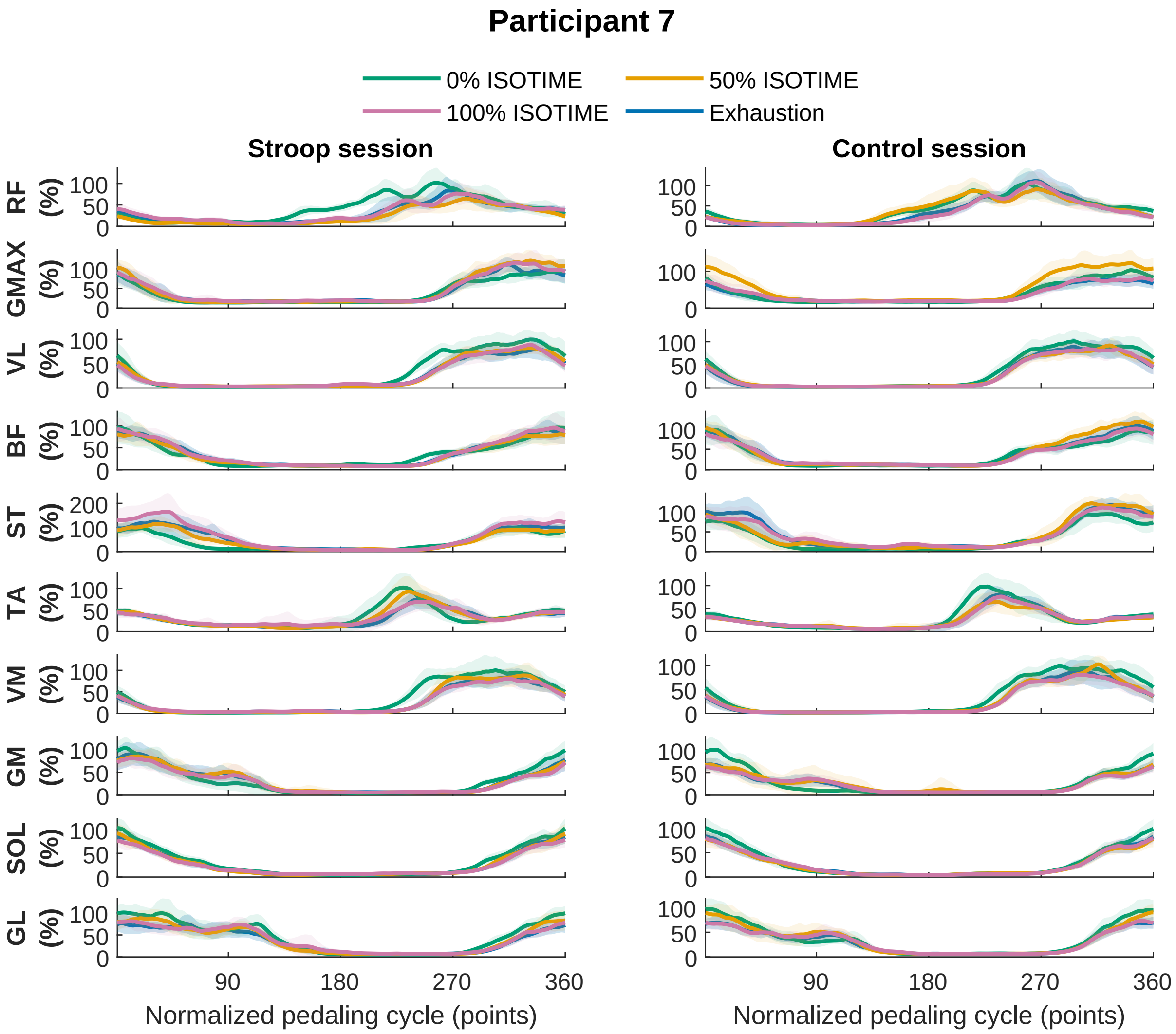

Participant 7
0% ISOTIME
50% ISOTIME
100% ISOTIME
Exhaustion
Stroop session
Control session
RF (%)
GMAX (%)
VL (%)
BF (%)
ST (%)
TA (%)
VM (%)
GM (%)
SOL (%)
GL (%)
0
50
100
200
90
180
270
360
Normalized pedaling cycle (points)

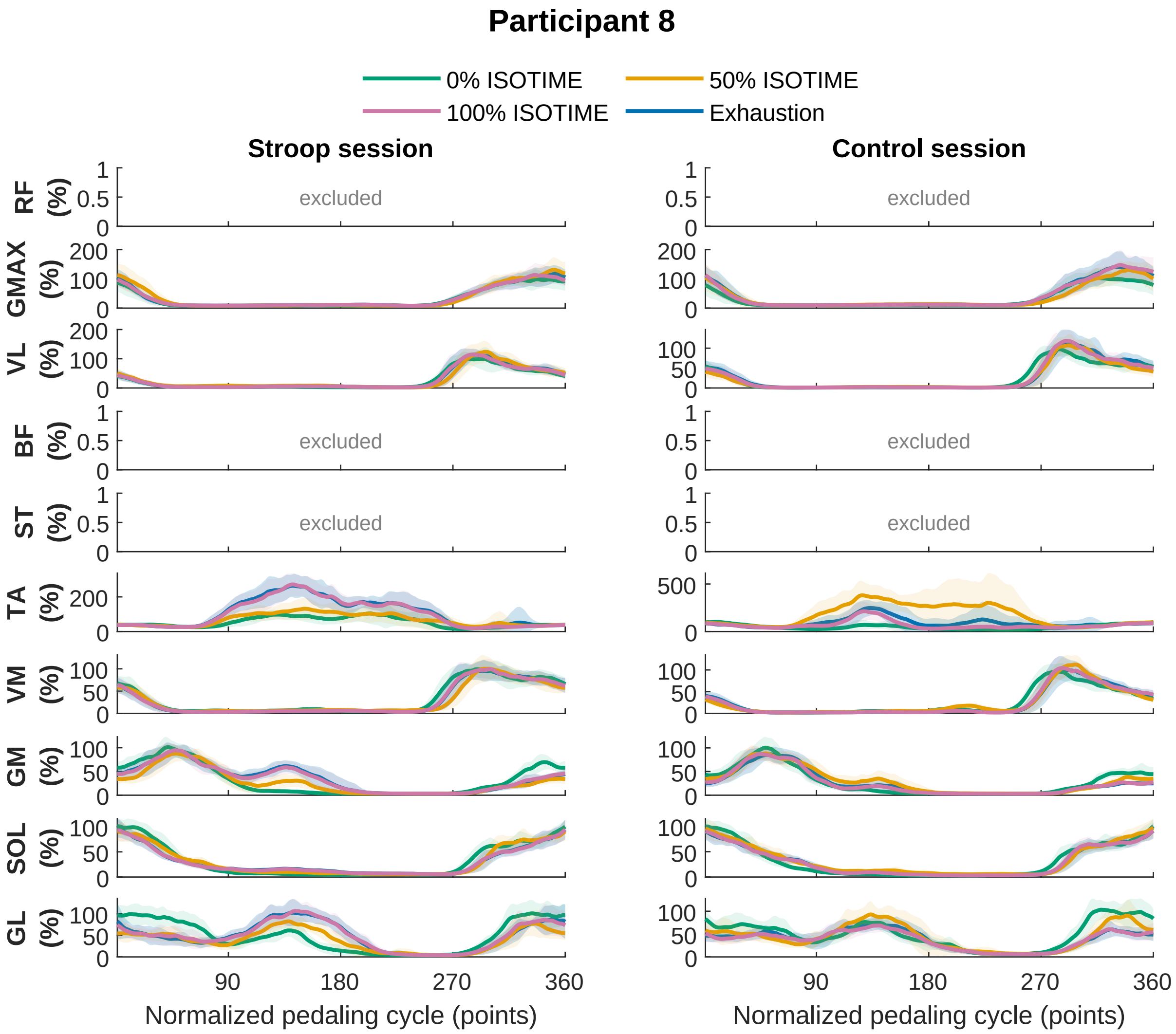
Participant 8
0% ISOTIME
50% ISOTIME
100% ISOTIME
Exhaustion
Stroop session
Control session
RF (%)
GMAX (%)
VL (%)
BF (%)
ST (%)
TA (%)
VM (%)
GM (%)
SOL (%)
GL (%)
excluded
90
180
270
360
Normalized pedaling cycle (points)

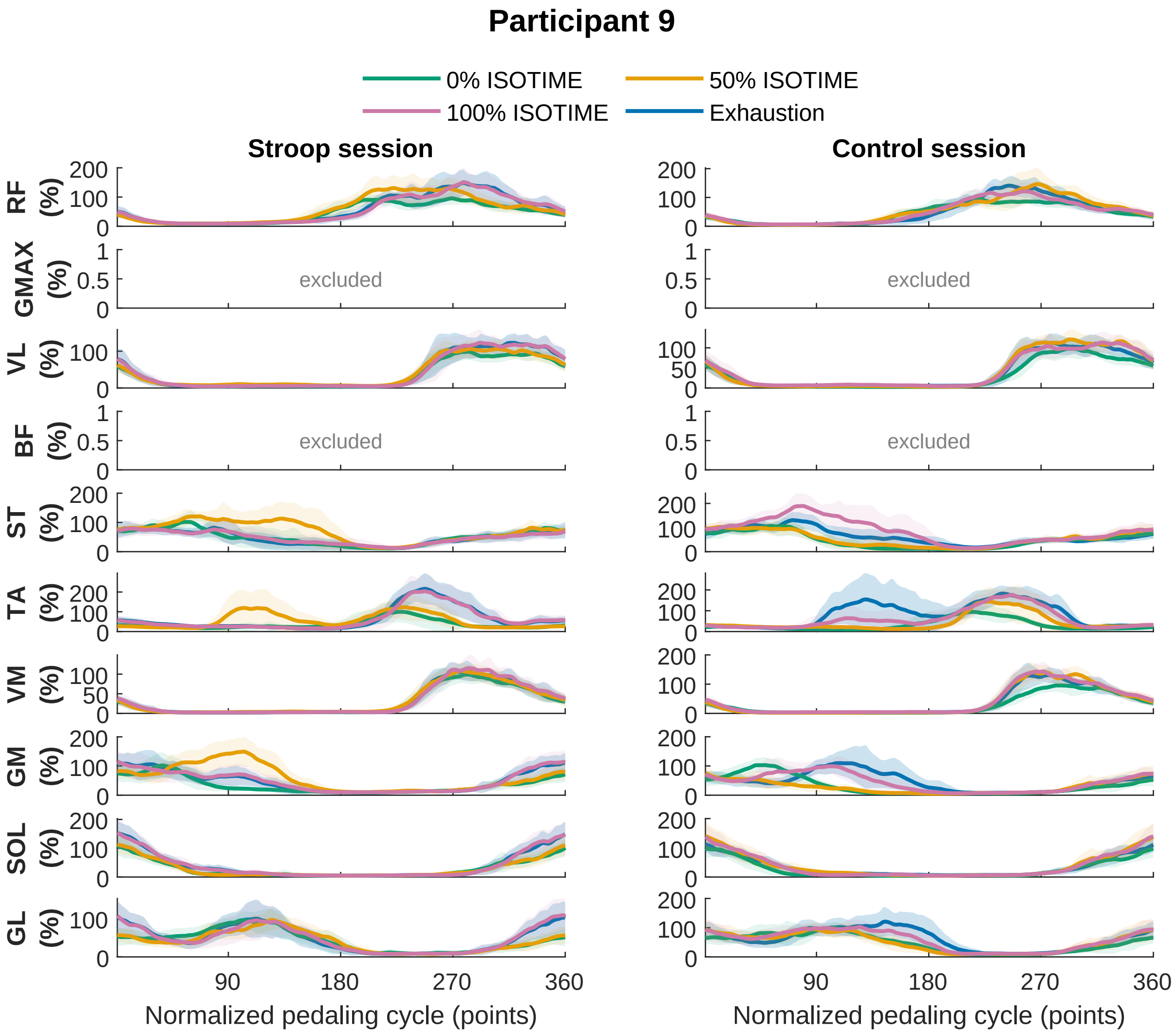

Participant 9
0% ISOTIME
50% ISOTIME
100% ISOTIME
Exhaustion
Stroop session
Control session
RF (%)
GMAX (%)
VL (%)
BF (%)
ST (%)
TA (%)
VM (%)
GM (%)
SOL (%)
GL (%)
excluded
Normalized pedaling cycle (points)

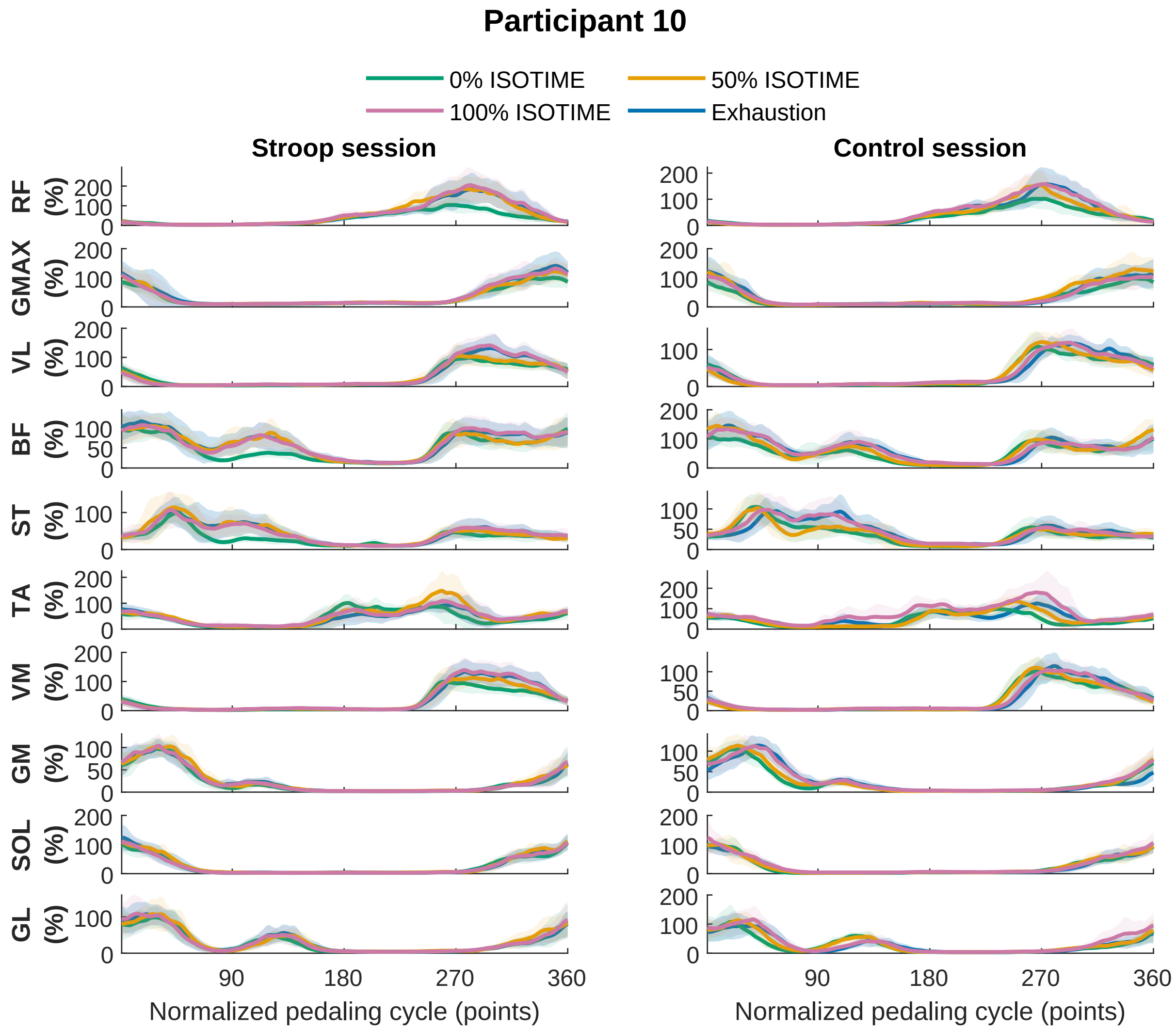
Participant 10
0% ISOTIME
50% ISOTIME
100% ISOTIME
Exhaustion
Stroop session
Control session
RF (%)
GMAX (%)
VL (%)
BF (%)
ST (%)
TA (%)
VM (%)
GM (%)
SOL (%)
GL (%)
90
180
270
360
Normalized pedaling cycle (points)
Normalized pedaling cycle (points)

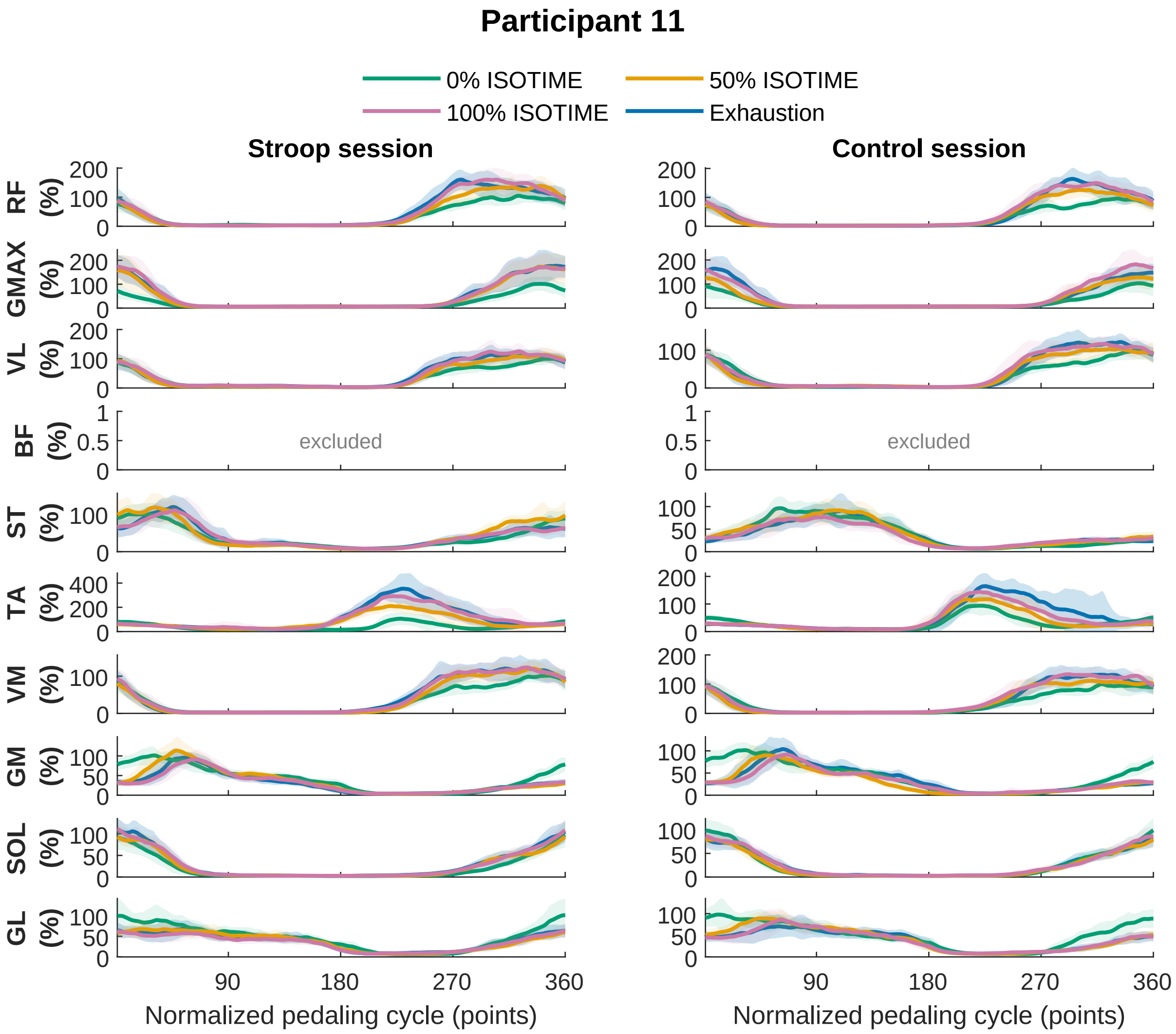
Participant 11
0% ISOTIME
50% ISOTIME
100% ISOTIME
Exhaustion
Stroop session
Control session
RF (%)
GMAX (%)
VL (%)
BF (%)
ST (%)
TA (%)
VM (%)
GM (%)
SOL (%)
GL (%)
excluded
excluded
90
180
270
360
Normalized pedaling cycle (points)
Normalized pedaling cycle (points)

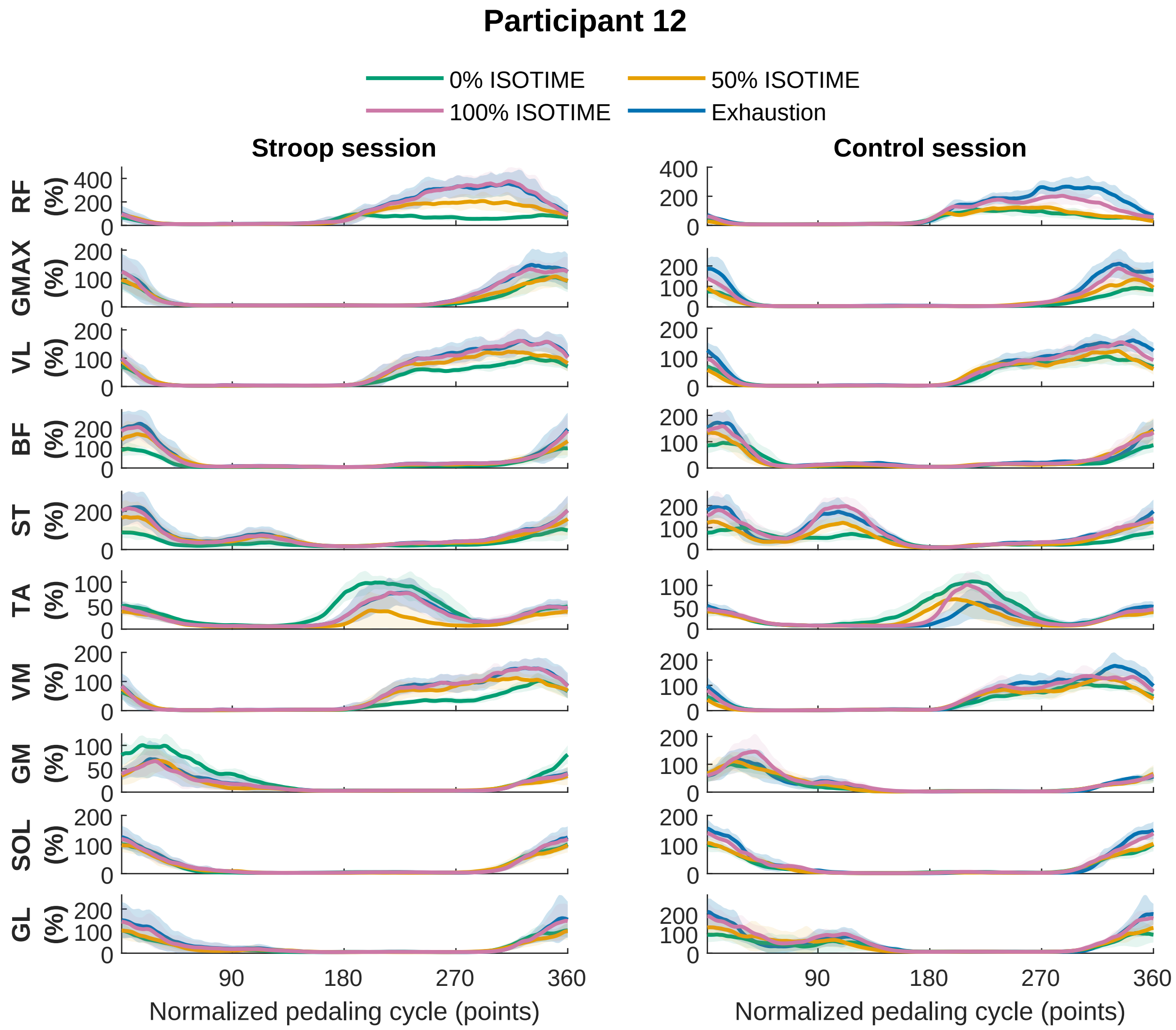
Participant 12
0% ISOTIME
50% ISOTIME
100% ISOTIME
Exhaustion
Stroop session
Control session
RF (%)
GMAX (%)
VL (%)
BF (%)
ST (%)
TA (%)
VM (%)
GM (%)
SOL (%)
GL (%)
Normalized pedaling cycle (points)
Normalized pedaling cycle (points)

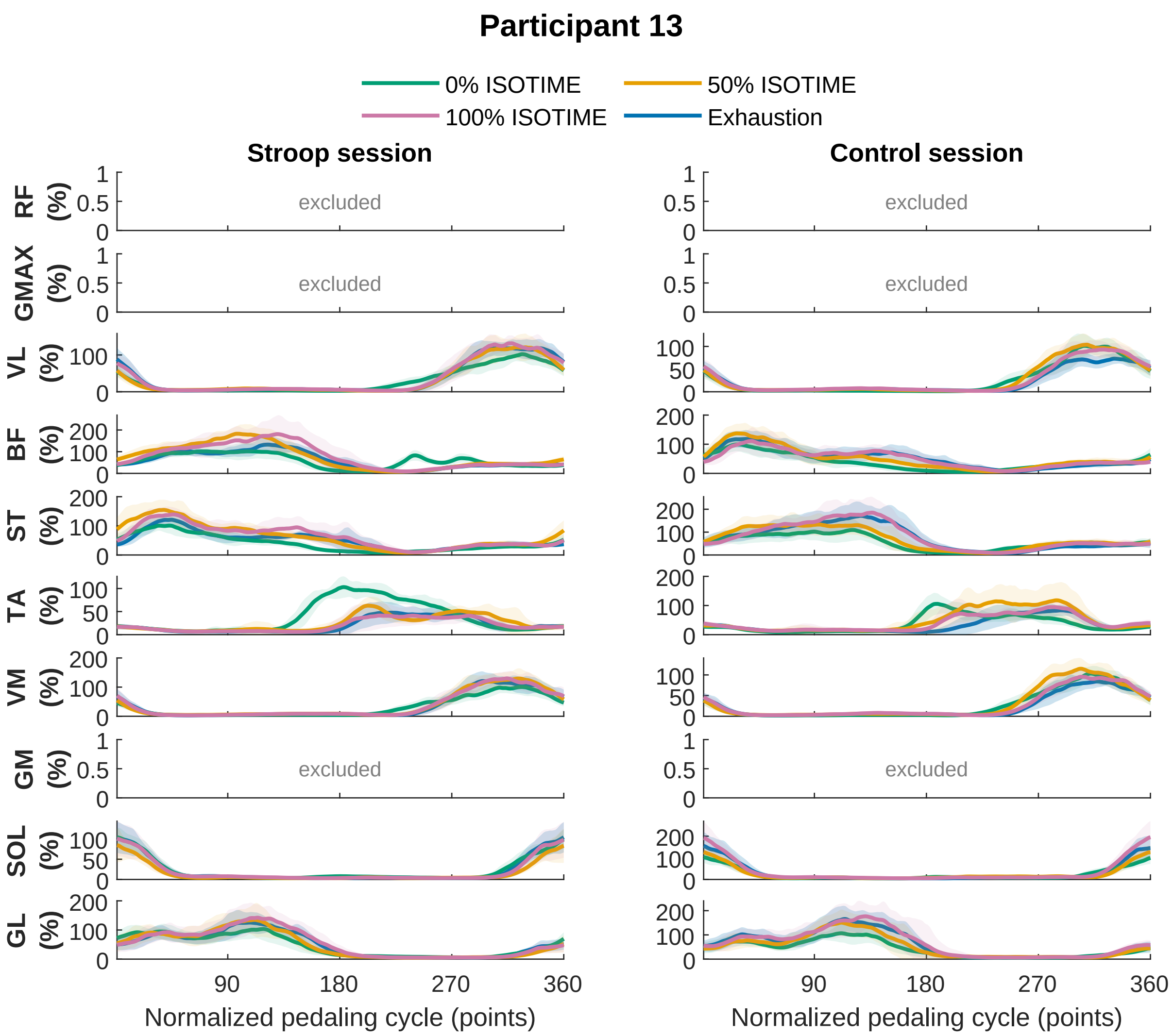

Participant 13
0% ISOTIME
50% ISOTIME
100% ISOTIME
Exhaustion
Stroop session
Control session
RF (%)
GMAX (%)
VL (%)
BF (%)
ST (%)
TA (%)
VM (%)
GM (%)
SOL (%)
GL (%)
excluded
90
180
270
360
Normalized pedaling cycle (points)

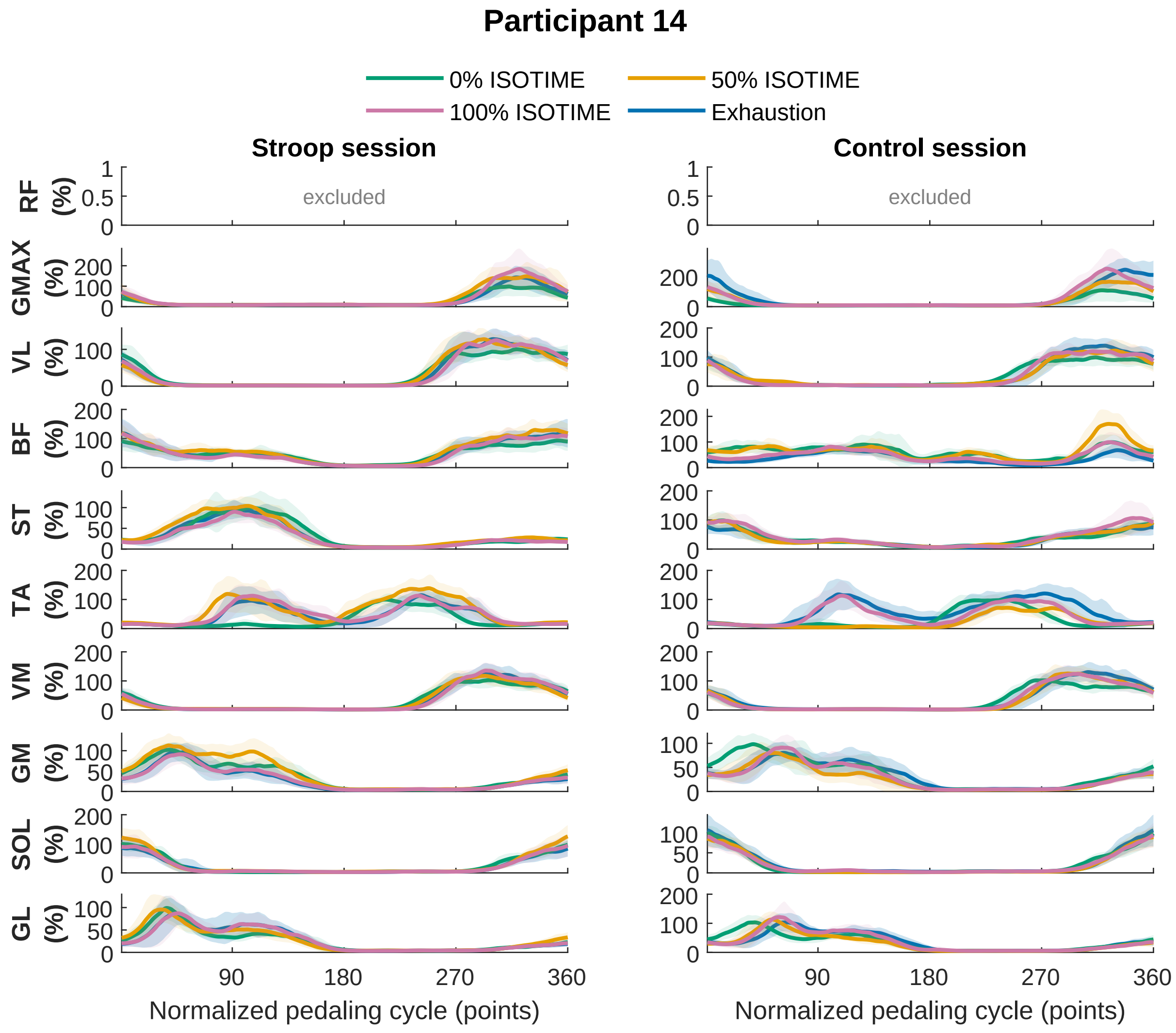
Participant 14
0% ISOTIME
50% ISOTIME
100% ISOTIME
Exhaustion
Stroop session
Control session
excluded
excluded
RF (%)
GMAX (%)
VL (%)
BF (%)
ST (%)
TA (%)
VM (%)
GM (%)
SOL (%)
GL (%)
Normalized pedaling cycle (points)
Normalized pedaling cycle (points)

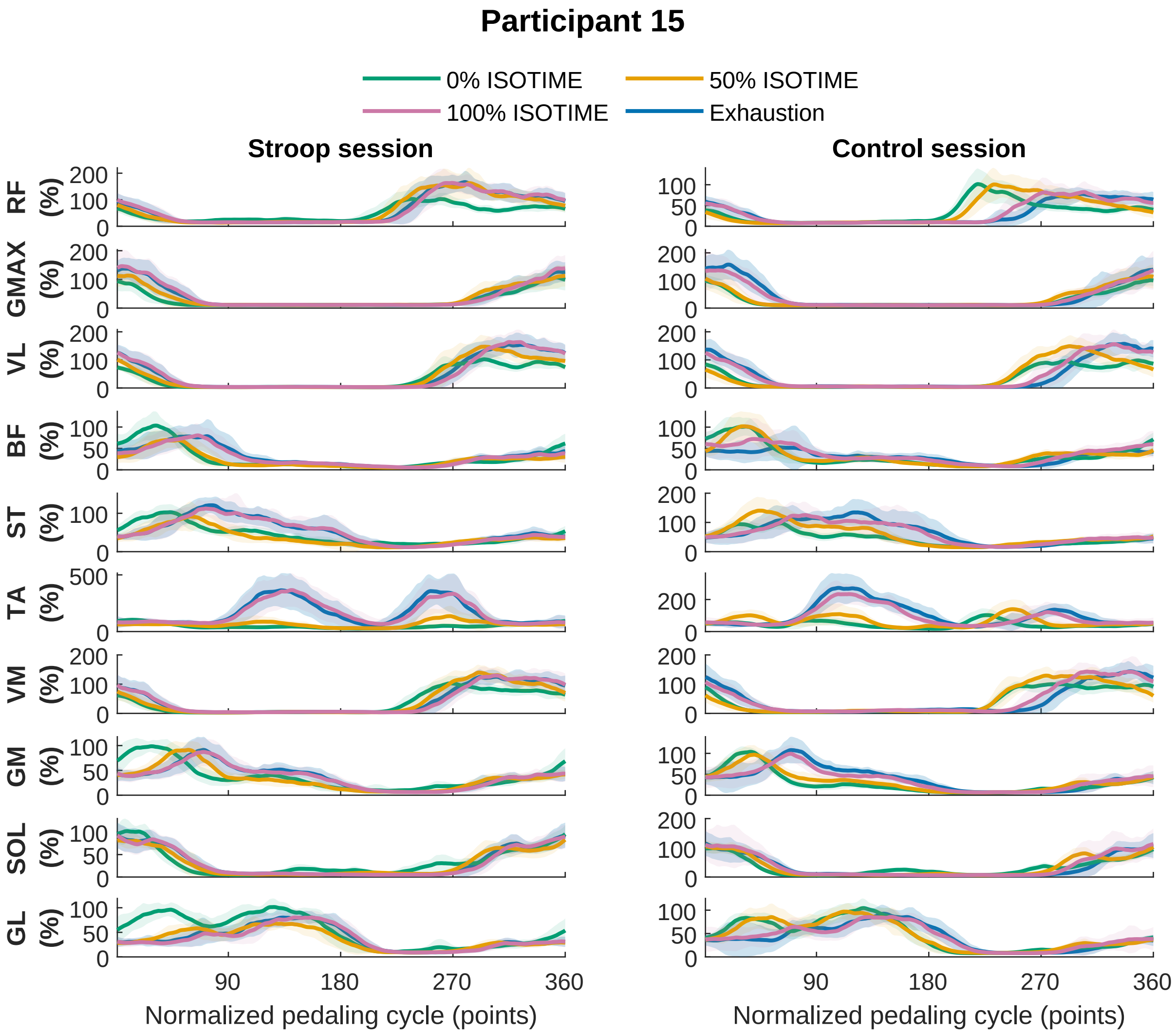

Participant 15
0% ISOTIME
50% ISOTIME
100% ISOTIME
Exhaustion
Stroop session
Control session
RF (%)
GMAX (%)
VL (%)
BF (%)
ST (%)
TA (%)
VM (%)
GM (%)
SOL (%)
GL (%)
90
180
270
360
Normalized pedaling cycle (points)

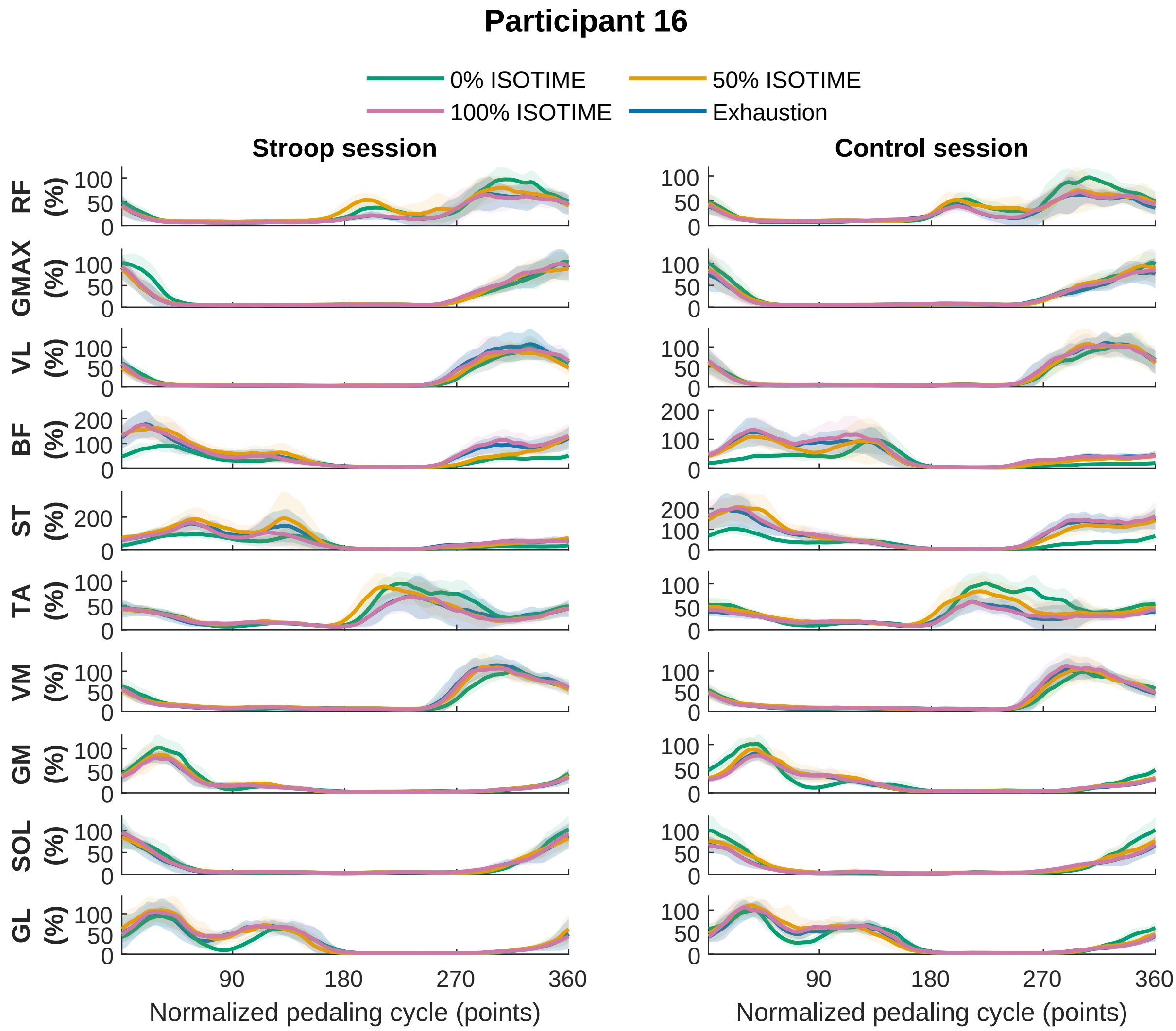
Participant 16
0% ISOTIME
50% ISOTIME
100% ISOTIME
Exhaustion
Stroop session
Control session
RF (%)
GMAX (%)
VL (%)
BF (%)
ST (%)
TA (%)
VM (%)
GM (%)
SOL (%)
GL (%)
90
180
270
360
Normalized pedaling cycle (points)
Normalized pedaling cycle (points)

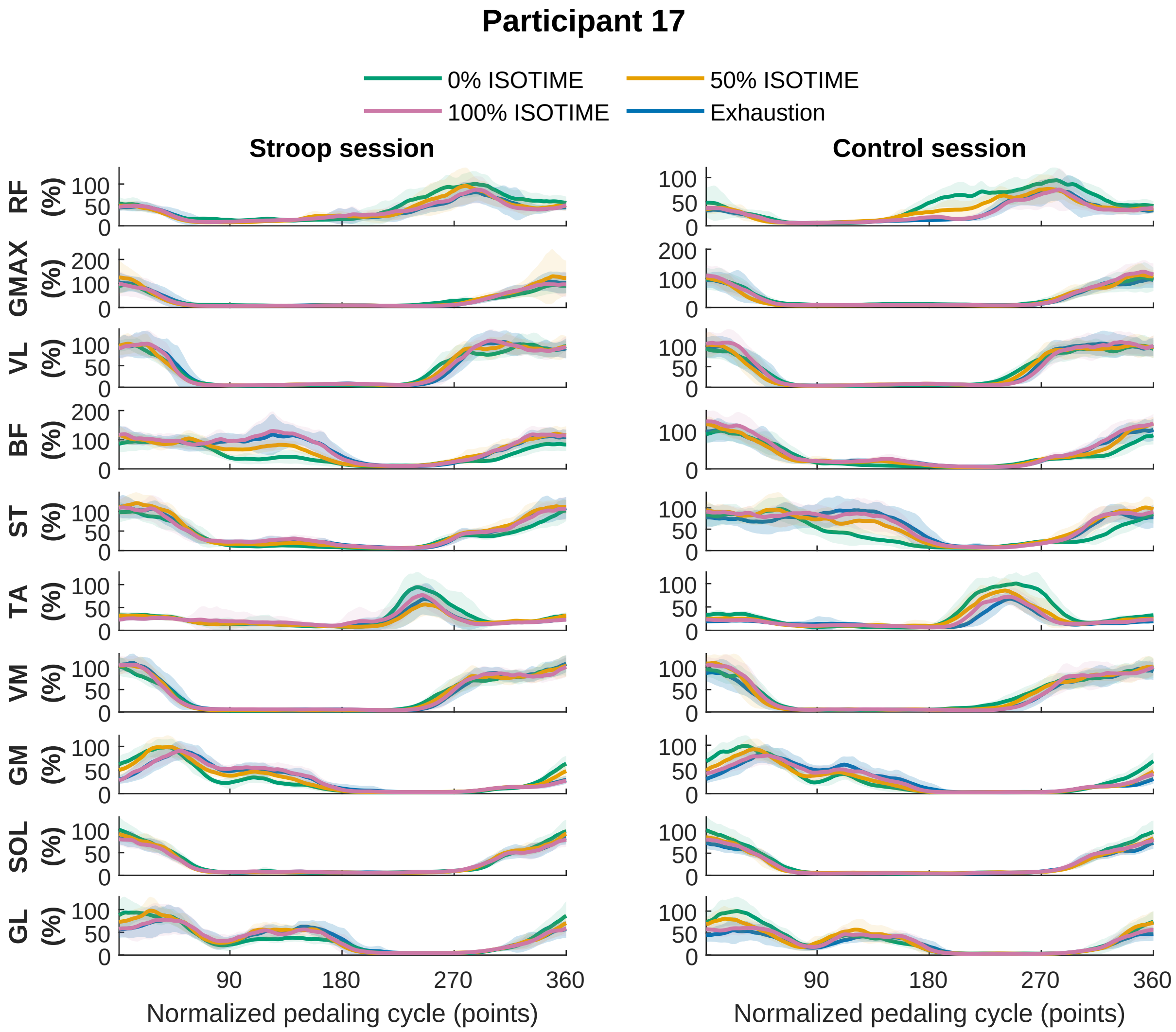
Participant 17
0% ISOTIME
50% ISOTIME
100% ISOTIME
Exhaustion
Stroop session
Control session
RF (%)
GMAX (%)
VL (%)
BF (%)
ST (%)
TA (%)
VM (%)
GM (%)
SOL (%)
GL (%)
90
180
270
360
Normalized pedaling cycle (points)
Normalized pedaling cycle (points)
