## Supplementary material for "Mental fatigue impairs cycling endurance performance and perception of effort, but not muscle activation": S3

**Supplementary Material S3. SPM1D results for EMG activity variability during the cycling time-to-exhaustion test (isotime analysis)**

| **Muscle** | **Tested effect** | **SPM statistic** | **df** | **p (corrected)** | **Significant cluster and cycle phase (%)** |
| --- | --- | --- | --- | --- | --- |
| RF | Session | F | 1,13 | > 0.05 (z_max = 8.20, z_threshold = 17.11) | – |
|  | Isotime | F | 2,26 | <0.0001 | [269 – 294] |
|  | Session × Isotime | F | 2,26 | > 0.05 (z_max = 5.53, z_threshold = 8.75) | – |
| GMax | Session | F | 1,13 | > 0.05 (z_max = 9.46, z_threshold = 17.20) | – |
|  | Isotime | F | 2,26 | 0.0123; 0.05 | [292 – 304]; [359 – 359] |
|  | Session × Isotime | F | 2,26 | > 0.05 (z_max = 3.37, z_threshold = 8.78) | – |
| VL | Session | F | 1,14 | > 0.05 (z_max = 6.67, z_threshold = 16.94) | – |
|  | Isotime | F | 2,28 | < 0.0001; 0.0004 | [263 – 293]; [302 – 322] |
|  | Session × Isotime | F | 2,28 | > 0.05 (z_max = 4.21, z_threshold = 8.76) | – |
| BF | Session | F | 1,12 | > 0.05 (z_max = 6.23, z_threshold = 17.57) | – |
|  | Isotime | F | 2,24 | 0.0495; 0.0310; 0.0142; 0.0488; 0.0259 | [28 – 29]; [43 – 51]; [304 – 317]; [326 – 328]; [331 – 340] |
|  | Session × Isotime | F | 2,24 | > 0.05 (z_max = 5.18, z_threshold = 8.82) | – |
| ST | Session | F | 1,15 | > 0.05 (z_max = 13.47, z_threshold = 15.76) | – |
|  | Isotime | F | 2,30 | 0.0483; 0.0030 | [91 – 93]; [319 – 336] |
|  | Session × Isotime | F | 2,30 | > 0.05 (z_max = 2.68, z_threshold = 8.38) | – |
| TA | Session | F | 1,16 | > 0.05 (z_max = 2.38, z_threshold = 14.39) | – |
|  | Isotime | F | 2,32 | 0.0340 | [292 – 300] |
|  | Session × Isotime | F | 2,32 | > 0.05 (z_max = 3.39, z_threshold = 7.88) | – |
| VM | Session | F | 1,16 | > 0.05 (z_max = 9.80, z_threshold = 15.78) | – |
|  | Isotime | F | 2,32 | 0.0117; 0.0342; 0.0349 | [106 – 117]; [131 – 137]; [269 – 274] |
|  | Session × Isotime | F | 2,32 | > 0.05 (z_max = 6.65, z_threshold = 8.44) | – |
| GM | Session | F | 1,13 | > 0.05 (z_max = 2.13, z_threshold = 17.45) | – |
|  | Isotime | F | 2,26 | > 0.05 (z_max = 6.89, z_threshold = 8.87) | – |
|  | Session × Isotime | F | 2,26 | > 0.05 (z_max = 2.26, z_threshold = 8.87) | – |
| SOL | Session | F | 1,16 | > 0.05 (z_max = 4.43, z_threshold = 15.65) | – |
|  | Isotime | F | 2,32 | > 0.05 (z_max = 5.11, z_threshold = 8.39) | – |
|  | Session × Isotime | F | 2,32 | > 0.05 (z_max = 4.34, z_threshold = 8.39) | – |
| GL | Session | F | 1,16 | > 0.05 (z_max = 9.35, z_threshold = 15.87) | – |
|  | Isotime | F | 2,32 | > 0.05 (z_max = 5.31, z_threshold = 8.48) | – |
|  | Session × Isotime | F | 2,32 | > 0.05 (z_max = 4.46, z_threshold = 8.48) | – |

Statistical Parametric Mapping (SPM1D) two-way repeated-measures ANOVAs were performed on normalized surface electromyography (EMG) signals variability over the full pedaling cycle (0–360%). Session (Stroop vs. control) and Isotime (0%, 50%, and 100% of individual isotime) were entered as within-subject factors. For each muscle, F-statistics, degrees of freedom (df), and Random Field Theory–corrected *p*-values are reported. When present, significant supra-threshold clusters (*p* < 0.05, corrected) are indicated together with the corresponding phase(s) of the pedaling cycle (%). For non-significant effects, the maximum test statistic (z_max) and the critical threshold (z_threshold) from SPM1D are reported, as exact *p*-values are not provided when no supra-threshold cluster is detected.

RF, rectus femoris; GMax, gluteus maximus; VL, vastus lateralis; BF, biceps femoris; ST, semitendinosus; TA, tibialis anterior; VM, vastus medialis; GM, gastrocnemius medialis; SOL, soleus; GL, gastrocnemius lateralis.
