## Supplementary material for "Mental fatigue impairs cycling endurance performance and perception of effort, but not muscle activation": S5

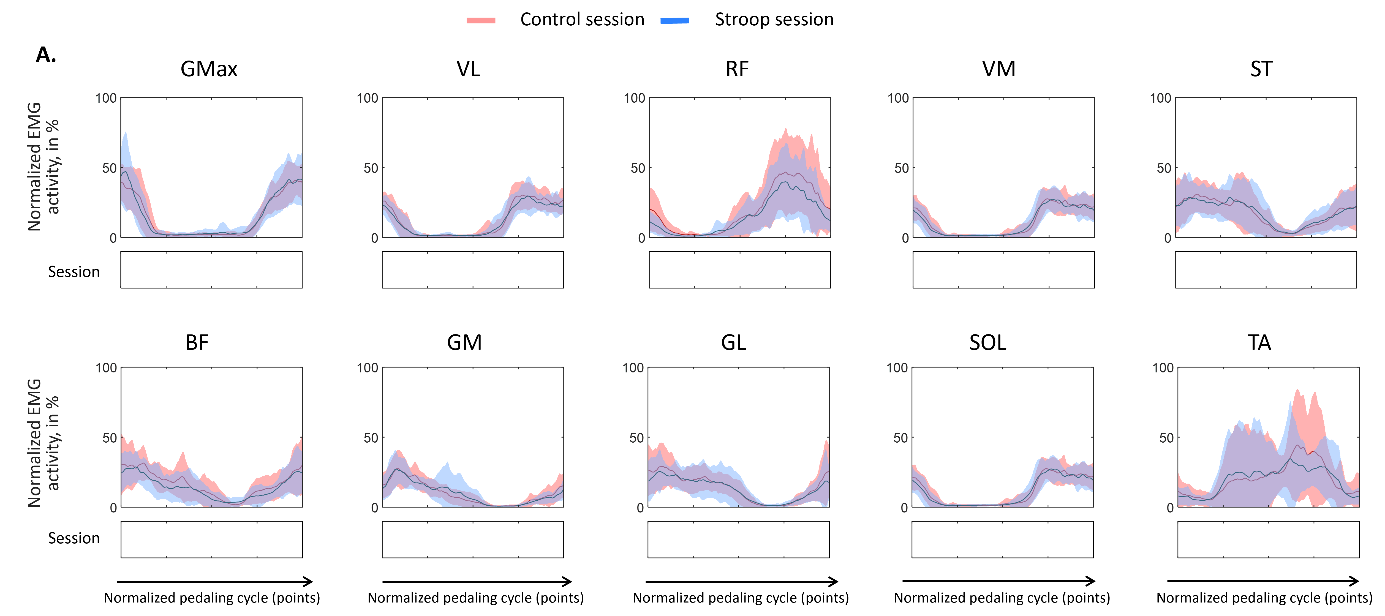

Supplementary material S5. **Variability (standard deviation) of electromyographic (EMG) activity of the lower limb muscles at exhaustion for the control (documentary) and Stroop conditions**. Standard deviation of EMG activity of the rectus femoris (RF), biceps femoris (BF), vastus medialis (VM), soleus (SOL), gastrocnemius lateralis (GL), gluteus maximus (GMax), vastus lateralis (VL), semi tendinous (ST), gastrocnemius medialis (GM) and tibialis anterior (TA) muscles throughout the pedaling cycle. EMG activity was measured at three isotime points (0%, 50%, and 100% – from top to bottom). Data are presented for two conditions: control (solid red line) and Stroop (solid blue line). The x-axis represents the normalized pedaling cycle (points), while the y-axis indicates normalized EMG activity. Solid lines represent the average muscle activity, with shaded areas denoting the standard deviation. Significant differences (*p* < 0.05) over time, assessed using a two-way repeated-measures ANOVA with factors task condition (Stroop vs. Control) and isotime (0%, 50%, and 100%), combined with Statistical Parametric Mapping analysis, are indicated by vertical black bars beneath each graph.
